# The engulfment receptor CED-1/MEGF10 activates the GPCR LAT-1/ADGRL3 for apoptotic cell degradation

**DOI:** 10.64898/2026.06.09.731128

**Authors:** Fuqiao Liu, Lei Yuan, Qian Zheng, Yang Kang, Aowei Wang, Kuo Li, Yunmin Xie, Lu Chen, Peiyao Li, Hui Wang, Zhi Li, Hui Xiao

## Abstract

Efferocytosis, the clearance of apoptotic cells, is crucial for tissue homeostasis, inflammation suppression, and repair. While several G protein-coupled receptors (GPCRs) are involved, the broader role of GPCRs in efferocytosis remains unexplored. Through a genome-wide RNAi screen in *Caenorhabditis elegans*, we identified the adhesion GPCR LAT-1 as a key regulator of apoptotic cell degradation through the promotion of phagosome maturation. Our secondary RNAi scrween for transcriptional regulators revealed that the transcription factor AST-1 acts downstream of LAT-1 to mediate apoptotic cell degradation. We show that the engulfment receptor CED-1 functions as a ligand for LAT-1, activating a Gs protein/adenylyl cyclase/PKA/AST-1 signaling cascade and inducing the transcription of *vps-34* and *piki-1*, which encode phosphatidylinositol 3-kinases essential for PtdIns3P generation on phagosomes. This pathway maintains appropriate VPS-34 and PIKI-1 levels for efficient efferocytosis. Notably, LAT-1 is evolutionarily conserved, with homologs in *Drosophila* (CIRL) and mammals (ADGRL3) performing similar functions. These findings reveal a conserved CED-1/MEGF10–LAT-1/ADGRL3 axis that orchestrates apoptotic cell clearance via a novel transcriptional regulatory pathway, providing new insight into the molecular mechanisms underlying tissue homeostasis and diseases associated with defective efferocytosis, including autoimmunity and neurodegeneration.

## Introduction

Programmed cell death through apoptosis is a fundamental biological process, eliminating approximately 200 billion cells daily in healthy human tissues^1^. Efferocytosis—the efficient clearance of apoptotic cells (ACs)—is mediated by professional phagocytes such as macrophages and dendritic cells, as well as by nonprofessional tissue-resident neighboring cells^2,3^. This process is essential for maintaining tissue homeostasis, suppressing inflammation, and promoting tissue repair. Defective efferocytosis contributes to the onset and progression of multiple autoimmune and inflammatory disorders, including systemic lupus erythematosus, atherosclerosis, and neurodegenerative diseases^4^. Efferocytosis is an evolutionarily conserved and tightly regulated process that is essential for normal development and tissue homeostasis^5,6^. The evolutionary conservation of this process across metazoans has enabled detailed mechanistic dissection using genetically tractable model organisms such as *Caenorhabditis elegans* and *Drosophila melanogaster*^7–9^.

In *C. elegans* hermaphrodites, 131 somatic cells and 300–500 germ cells undergo apoptosis and are subsequently phagocytosed and degraded by neighboring somatic or gonadal sheath cells^10^. These apoptotic cells, referred to as cell corpses, appear as distinct “button-like” structures under differential interference contrast (DIC) microscopy^11^. Defects in corpse clearance lead to the persistence and accumulation of apoptotic cells, resulting in a phenotype termed Ced (cell death abnormal). During apoptosis in *C. elegans*, the activation of CED-8 by the caspase CED-3 triggers the exposure of phosphatidylserine (PS) on the dying cell surface as an “eat-me” signal^12,13^ This signal is recognized by phagocytes through two partially redundant genetic pathways. The *ced-2/5/12* pathway activates the small GTPase CED-10/Rac1, driving actin cytoskeletal rearrangement for cell corpse engulfment^14,15^. In contrast, the *ced-1/6/7* pathway has dual functions—regulating both engulfment and degradation^16,17^. Within this pathway, TTR-52 binds exposed PS and the extracellular domain of CED-1, acting as a bridging molecule that links dying cells to phagocytes^18^. CED-7, a mammalian ABC transporter homolog, functions in both dying and engulfing cells and is essential for CED-1 enrichment around cell corpses^19^. The phagocytic receptor CED-1 recognizes PS on apoptotic cells and clusters on the plasma membrane adjacent to the corpse. This clustering promotes pseudopod extension and closure of the phagocytic cup to form a phagosome^20^. Beyond engulfment, CED-1 also initiates degradation^21^. It does so by recruiting class II PI3K (PIKI-1), class III PI3K (VPS-34), and the small GTPases RAB-5 and RAB-7 to nascent phagosomes through CED-6 and DYN-1^22^. Rab GTPases (RAB-5, −7, −14, and UNC-108/RAB-2) and their regulatory effectors act sequentially to drive phagosome maturation, culminating in phagolysosome formation and lysosomal degradation of cell corpses^23^. While the molecular mechanisms of efferocytosis, particularly in model organisms such as *C. elegans*, have been extensively studied, the involvement of major cell surface receptor families, such as G protein-coupled receptors (GPCRs), in this process remains unclear.

G protein–coupled receptors (GPCRs) constitute the largest family of cell-surface receptors; these receptors are conserved across a wide range of organisms and are integral to numerous physiological and pathological processes^24,25^. More than 800 GPCRs have been identified in the human genome and are grouped into five major classes on the basis of sequence similarity and evolutionary origin^26^. Despite sharing a canonical seven-transmembrane (7TM) architecture, GPCRs exhibit extensive diversity in ligand binding, downstream signaling, and biological functions^26^. Identifying physiologically relevant ligands for many GPCRs remains a key challenge. Despite the diverse roles of GPCRs in cellular processes and the fact that a few specific receptors, such as BAI1, GPR132, and GPR37, have been implicated in efferocytosis^27–29^, a systematic investigation of the involvement of GPCRs in efferocytosis is lacking, and the mechanisms through which GPCRs coordinate with the core engulfment machinery remain unknown.

Given the critical role of efficient efferocytosis in preventing autoimmune and inflammatory disorders and the therapeutic potential of targeting GPCRs, there is an unmet need to systematically identify and characterize novel GPCRs that regulate distinct phases of apoptotic cell clearance. To address this knowledge gap and identify novel GPCR regulators of efferocytosis, we conducted a genome-wide RNAi screen in *Caenorhabditis elegans*. Our screen pinpointed the adhesion GPCR LAT-1 (mammalian ortholog ADGRL3/latrophilin-3) as a critical and previously unappreciated player in apoptotic cell clearance. Through a secondary genome-wide screen for transcriptional regulators, we discovered that LAT-1 activates a G protein/adenylyl cyclase/PKA signaling cascade that induces the expression of the PI3Ks VPS-34 and PIKI-1 via the transcription factor AST-1 (mammalian ortholog ETV1). We further demonstrated that the engulfment receptor CED-1 physically interacts with LAT-1 to coordinate apoptotic cell recognition with transcriptional upregulation of the phagosome maturation machinery. Importantly, we report that this CED-1/MEGF10–LAT-1/ADGRL3 axis is evolutionarily conserved, with mammalian ADGRL3 performing analogous functions in macrophage-mediated efferocytosis. Our findings reveal an unexpected function for LAT-1/ADGRL3 in apoptotic cell clearance and provide insights into how adhesion-GPCR signaling integrates with canonical efferocytosis pathways, highlighting LAT-1/ADGRL3 as a potential therapeutic target for diseases linked to defective efferocytosis and dysregulated apoptosis.

## Results

### Identification and validation of LAT-1 as a regulator of apoptotic cell clearance in *C. elegans*

To identify GPCRs potentially involved in apoptotic cell clearance, we individually knocked down 709 GPCR genes (WormBase: G protein-coupled receptor signaling pathway, GO: 0007186; Table EV1) and screened for germline cell death across three independent replicates. Inactivation of 42 GPCR genes increased the number of germline cell corpses in at least two experiments (Table EV2).

To determine whether this increase resulted from enhanced apoptosis or defective clearance, we performed RNAi in *xwhIs49 (P_ced-1_rde-1; rde-1)* worms (for phagocyte-specific RNAi suppression) and *xwhIs82 (P_egl-1_rde-1; rde-1)* worms (for apoptotic cell-specific RNAi suppression). RNAi knockdown of five GPCR genes—*srw-40*, *lat-1*, *seb-3*, *sra-11*, and *W08G11.5*—significantly increased embryonic cell corpses in *P_ced-1_rde-1; rde-1* worms but not in *P_egl-1_rde-1; rde-1* worms compared with their respective controls (Table EV3–4). These findings indicate that silencing these genes impaired apoptotic cell clearance rather than inducing apoptosis. Among them, *lat-1* knockdown caused the most pronounced accumulation of cell corpses at multiple embryonic stages, prompting its selection for further analysis (Fig. 1a).

**Fig. 1.**
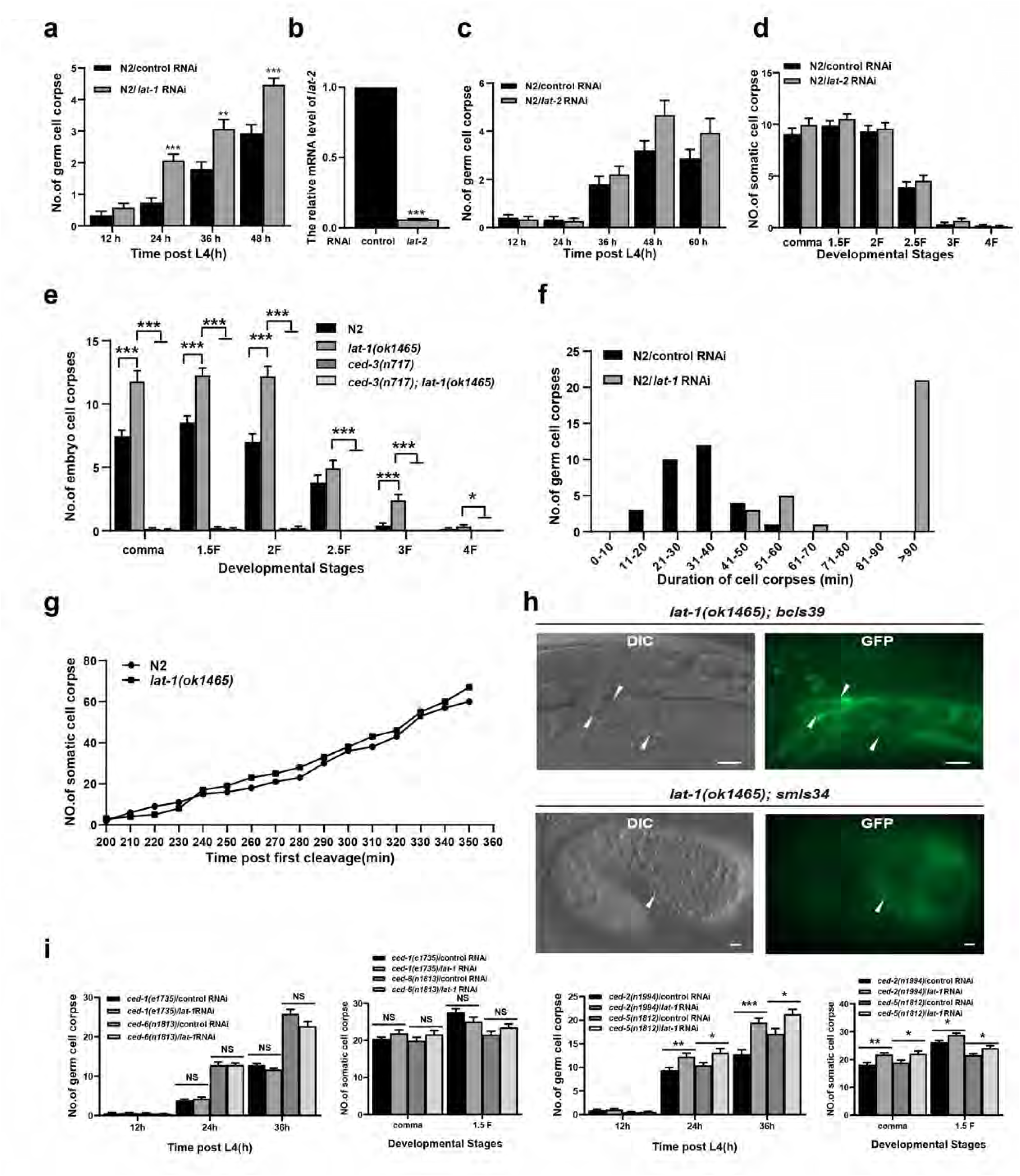
LAT-1 regulates cell corpse clearance in *C. elegans*. (a, c-e, and i) Quantification (mean ± SEM) of button-like cell corpses at different adult stages (h post L4) in the germline or at different embryonic stages in the soma of the indicated strains. Fifteen adult worms or fifteen embryos were scored at each stage for each strain. (b) The relative mRNA level o*f lat-2* was detected by qPCR. (f) Four-dimensional microscopy analysis of germ cell corpse duration in N2/control and *lat-1* RNAi worms. The persistence of 30 germ cell corpses was also monitored. (g) The number of embryo corpses was measured using four-dimensional microscopy in N2/control and *lat-1* RNAi worms. Embryos (2-cell stage) were isolated from adult worms and imaged every minute for 400 min. (h) CED-1::GFP localization in germlines (top, *lat-1(ok1465); bcIs39* strain) and embryos (bottom, *lat-1(ok1465)*; *smIs34* strain) was imaged by the GFP channel using an Imager M2 microscope. The arrows indicate cell corpses. Scale bars: 10 µm (top) and 2 µm (bottom). Unpaired t tests were performed. *P < 0.05, **P < 0.01, ***P < 0.001; NS, not significant. All error bars indicate the mean ± SEM.

*C. elegans* contains two latrophilin homologs, *lat-1* and *lat-2*, with *lat-1* being more closely related to latrophilin. *lat-1* is involved in anterior–posterior patterning during early embryogenesis and in planar cell polarity (PCP)^30,31^. *C. elegans* LAT-1 and LAT-2 share 21% amino acid identity. *lat-2* displays a more restricted expression pattern and increases the lethality of *lat-1* mutants^30^. Using qPCR, we confirmed that *lat-2* mRNA levels were efficiently reduced by RNAi (Fig. 1b). However, the number of cell corpses did not significantly differ between *lat-2* RNAi and control worms (Fig. 1c-d), indicating that *lat-2* does not participate in apoptotic cell clearance.

To further define the role of *lat-1*, we analyzed *lat-1(ok1465)* deletion mutants, which represent strong loss-of-function alleles^30^. Compared with the wild-type (WT) controls, these mutants resulted in significantly increased numbers of embryonic and germline cell corpses (Fig. 1e). Because *ced-3* and *ced-4* are essential for apoptotic cell death in *C. elegans*^32^, loss-of-function mutations in either gene almost completely abolish apoptosis. Accordingly, compared with the *lat-1(ok1465)* mutants alone, the *lat-1(ok1465); ced-3(n717)* double mutants displayed markedly fewer embryonic and germline cell corpses (Fig. 1e). Furthermore, both embryonic and germ cell corpses in *lat-1(ok1465)* mutants were labeled by the phagocytic receptor CED-1 (Fig. 1f), confirming that the accumulated corpses originated from apoptotic cell death.

We next used four-dimensional microscopy to measure the persistence of germ cell corpses. In WT animals, germ cell corpses rarely persisted for more than 60 min, whereas *lat-1* knockdown resulted in prolonged corpse persistence (>100 min) in most gonadal germ cells (Fig. 1g). However, the total number of cell deaths in *lat-1(ok1465)* embryos was indistinguishable from that in WT embryos (Fig. 1h). Together, these results indicate that the adhesion GPCR LAT-1 contributes to apoptotic cell clearance rather than the initiation of apoptosis.

Phagocytosis of apoptotic cells in *C. elegans* is controlled by two partially redundant signaling pathways: CED-1/CED-6/CED-7/NRF-5/TTR-52/DYN-1 and CED-2/CED-5/CED-10/CED-12/PSR-1^10^. To determine whether LAT-1 functions within one of these pathways, we treated the *ced-1(e1735)* and *ced-6(n1813)* engulfment mutants with *lat-1* RNAi. *lat-1* silencing did not further exacerbate the defects in germ cell engulfment in either mutant (Fig. 1i). In contrast, *lat-1* RNAi treatment markedly worsened engulfment defects in *ced-2(n1994)* and *ced-5(n1812)* mutants, which are components of the alternate pathway (Fig. 1i). These findings suggest that LAT-1 acts in the CED-1/6/7 genetic pathway to promote cell corpse removal.

To examine the defects in cell corpse clearance caused by the loss of *lat-1*, we assessed the phagosomal recruitment of factors involved in phagosome maturation. Persistent germ cell corpses in *lat-1* knockdown worms were surrounded by markers of engulfing cells, including CED-1::GFP, CED-6::GFP, DYN-1::GFP, APA-2::GFP, GFP::moesin, and ACT-1::mCherry (Fig. EV1a), confirming successful engulfment. We next analyzed the following GFP-tagged markers of distinct phagosome maturation stages: the early phagosome markers YFP::2xFYVE (a PtdIns3P biosensor) and GFP::RAB-5; the early maturation factor LST-4::GFP; the late endosome marker GFP::RAB-7; the lysosomal membrane proteins LMP-1::mCHERRY and LAAT-1::GFP; and the lysosomal hydrolases NUC-1::mCHERRY and CPL-1::mChOint. While the proportions of phagosomes surrounded by GFP::RAB-5, LST-4::GFP, GFP::RAB-7, LAAT-1::GFP, NUC-1::mCHERRY, and CPL-1::mChOint were comparable between lat-1 RNAi and control worms, significantly fewer phagosomes exhibited YFP::2xFYVE enrichment under *lat-1* knockdown, indicating impaired recruitment of PtdIns3P during early phagosome maturation.

A crucial step in phagosome maturation is the gradual acidification of the phagosomal lumen, as an acidic environment promotes the activity of hydrolytic enzymes that degrade phagosomal contents^33^. To test whether cell corpse persistence in *lat-1* RNAi and *lat-1* mutants reflected defective phagosomal acidification, a late stage of clearance, we stained worms with acridine orange (AO). AO-positive phagosomes were markedly reduced in both *lat-1* RNAi and *lat-1* mutants relative to those in controls (Fig. EV1b), supporting the notion that LAT-1 loss impairs phagosomal acidification following engulfment.

We next examined the effect of LAT-1 on endosomal–lysosomal degradation using *arIs36(Pheat-shockssGFP)* worms expressing secreted GFP (ssGFP) under a heat-shock promoter. In control worms, ssGFP accumulated in the body cavity within 1 h after heat shock (hphs), was efficiently taken up by coelomocytes within 12 hphs, and was completely degraded by 24 hph (Fig. EV1c). In *lat-1* knockdown worms, ssGFP uptake by coelomocytes occurred normally, but ssGFP persisted in both the body cavity and coelomocytes up to 48 hph (Fig. EV1c), indicating severely impaired endosomal/lysosomal degradation.

To directly visualize chromatin degradation during phagosome maturation, we used *ujIs113 (Ppie-1H2B::mCherry; Pnhr-2HIS-24::mCherry)* worms expressing germline-specific chromatin markers. In control worms, condensed chromatin in germ cell corpses disappeared within ∼30 min (31.0 ± 2.83 min, *n* = 5). In *lat-1* RNAi worms, the chromatin initially condensed but subsequently diffused throughout the corpse, with mCherry signals persisting for more than 70 min (80.0 ± 7.94 min, *n* = 5) (Fig. EV1d). These findings demonstrate that LAT-1 deficiency delays chromatin degradation within phagosomes, indicating a defect in postengulfment processing. Collectively, these results show that the accumulation of apoptotic cells in *lat-1*-knockdown worms results from defective degradation within engulfing cells.

### GPCR-dependent G protein signaling mediated by LAT-1 regulates apoptotic cell clearance

Upon activation, adhesion GPCRs (aGPCRs) can signal through canonical G protein-dependent or noncanonical G protein–independent pathways^34^. Canonical signaling involves heterotrimeric G proteins (Gαs, Gαi, Gα12/13, or Gαq), whereas noncanonical signaling involves the recruitment of monomeric intermediates such as β-arrestins, RacGEFs, or RhoGEFs^34^. Previous studies have shown that LAT-1 controls early embryonic cell division alignment in *C. elegans* via a metabotropic Gs/adenylyl cyclase/cAMP pathway^35^. We therefore hypothesized that LAT-1 regulates apoptotic cell clearance through a similar canonical signaling cascade.

The *C. elegans* genome encodes 21 Gα, 2 Gβ, and 2 Gγ subunits, as well as 4 adenylyl cyclases^36^. We individually knocked down each of these genes and screened for germline cell death in triplicate. Inactivation of *gpa-7*, *gpa-13*, and *acy-3* increased the number of germline cell corpses in at least two replicates (Table EV5). To determine whether these increases reflected enhanced apoptosis or defective clearance, we performed RNAi in *xwhIs49 (P_ced-1_rde-1; rde-1)* and *xwhIs82 (P_egl-1_rde-1; rde-1)* transgenic worms. Knockdown of *gpa-7* expression significantly increased embryonic cell corpses in *P_ced-1_rde-1; rde-1* worms in at least two replicates but not in *P_egl-1_rde-1; rde-1* worms, indicating a defect in corpse clearance rather than apoptosis induction (Table EV6–7). Knockdown of *gpa-13* slightly decreased the number of embryonic corpses in *P_ced-1_rde-1; rde-1* worms in at least two replicates, suggesting a potential negative regulatory role. The knockdown of *acy-3* expression significantly increased the number of embryonic corpses in both transgenic strains, suggesting that it plays a role in both processes. However, its clearance phenotype is more pronounced (Table EV6–7).

Because Gs directly activates adenylyl cyclase to generate cAMP, which in turn activates protein kinase A (PKA), we tested whether *kin-1*, the *C. elegans* gene encoding the catalytic subunit of PKA, mediates apoptotic cell clearance. RNAi knockdown of *kin-1* increased germline cell corpses and specifically enhanced embryonic corpses in *P_ced-1_rde-1; rde-1* worms but not in *P_egl-1_rde-1; rde-1* worms, suggesting a defect in corpse removal rather than apoptosis induction (Table EV6–7).

To assess genetic interactions, we treated *lat-1(ok1465)* mutants with *gpa-7*, *acy-3*, or *kin-1* RNAi. The knockdown of these genes did not further exacerbate the engulfment defects of *lat-1(ok1465)* mutants (Fig. 2a–c), indicating that they function in the same genetic pathway as *lat-1* does. Coimmunoprecipitation (Co-IP) assays revealed that LAT-1 interacts with GPA-7; in turn, GPA-7 interacts with ACY-3 (Fig. 2d-e). Yeast two-hybrid (Y2H) assays confirmed direct interactions between GPA-7 and both LAT-1 and ACY-3 (Fig. 2f-g). *gpa-7* is expressed in most neurons and all muscle cells, intestinal cells, and gonad sheath cells^36^, which are amateur cells engulfed for the removal of apoptotic cells in *C. elegans*. Although *gpa-7* is not a clear ortholog of mammalian proteins, our results indicate that it acts as a Gs-like protein in apoptotic cell clearance, implying functional similarity despite sequence divergence. Together, these findings support a model in which LAT-1 activates the Gs/adenylyl cyclase/cAMP/PKA signaling cascade to promote apoptotic cell clearance.

**Fig. 2.**
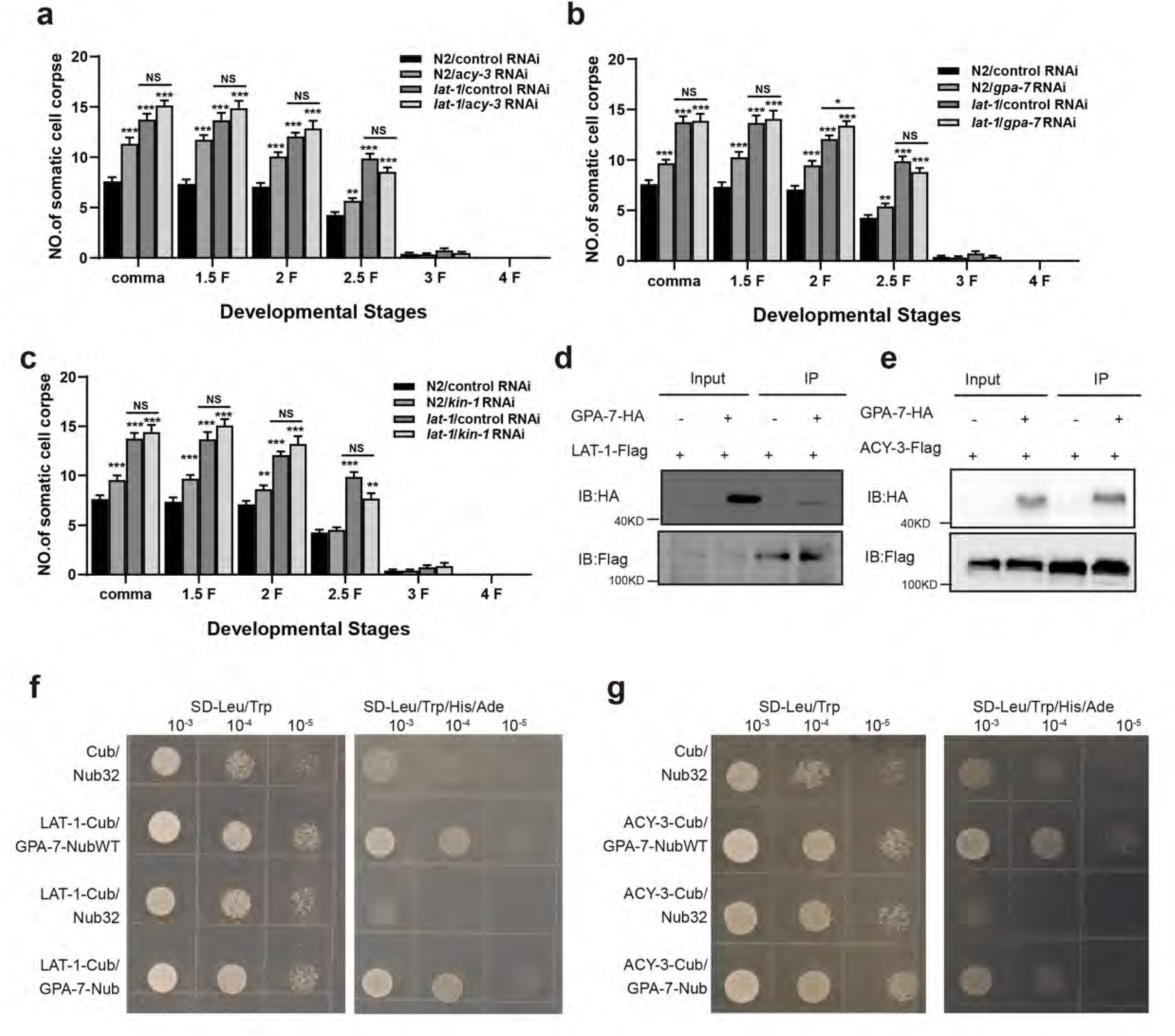
LAT-1 activates the GPA-7/ACY-3/KIN-1 pathway to promote cell corpse clearance. (a–c) Quantification (mean ± SEM) of embryonic cell corpses at different embryonic stages using the indicated strains. Fifteen embryos were scored at each embryonic stage for each strain. (d–e) The interactions between GPA-7 and LAT-1 (d) and between GPA-7 and ACY-3 (e) were examined by co-IP in 293T cells. (f and g) Yeast two-hybrid assay of the interactions between GPA-7 and LAT-1 (f) and between GPA-7 and ACY-3 (g). SD-Leu/Trp, medium lacking Leu and Trp; SD-Leu/Trp/His/Ade, medium lacking Leu, Trp, His, and Ade.

### The LAT-1 signaling pathway upregulates *vps-34* and *piki-1* transcription

We observed a significant reduction in the proportion of phagosomes enriched in YFP::2xFYVE following *lat-1* knockdown (Fig. EV1a). Because 2xFYVE is a PtdIns3P biosensor and PtdIns3P transiently accumulates on apoptotic cell–containing phagosomes at an early maturation stage under the control of class II PI3K PIKI-1 and class III PI3K VPS-34, these data suggest that PtdIns3P production is defective^37^. The PI 3-phosphatase MTM-1 antagonizes PIKI-1/VPS-34 by downregulating PtdIns(3)P on phagosomes; thus, MTM-1 coordinates with PIKI-1 to tune PtdIns3P levels^37^.

We found that knockdown of *vps-34* or *piki-1* significantly decreased the percentage of phagosomes surrounded by YFP::2xFYVE, which is consistent with the findings of previous studies (Fig. 3a)^38^. We next tested whether LAT-1 regulates the transcription of *vps-34* and *piki-1*. Quantitative PCR (qPCR) revealed decreased *vps-34* and *piki-1* mRNA in *lat-1* RNAi-treated worms and in *lat-1(ok1465)* mutants (Fig. 3b-c). The transcription of *vps-34* and *piki-1* was likewise reduced upon RNAi of *gpa-7*, *acy-3*, or *kin-1* (Fig. 3c-f). Consistent with these findings, knockdown of *gpa-7*, *acy-3*, or *kin-1* significantly decreased the fraction of YFP::2xFYVE-positive phagosomes, whereas the percentages of phagosomes surrounded by CED-1::GFP, GFP::RAB-5, GFP::RAB-7, LAAT-1::GFP, NUC-1::mCHERRY, or CPL-1::mChOint did not significantly differ from those of the vector controls (Fig. 3g). These results indicate that the LAT-1/Gs/adenylyl cyclase/PKA pathway promotes PtdIns3P enrichment on phagosomes by activating the transcription of VPS-34 and PIKI-1.

**Fig. 3.**
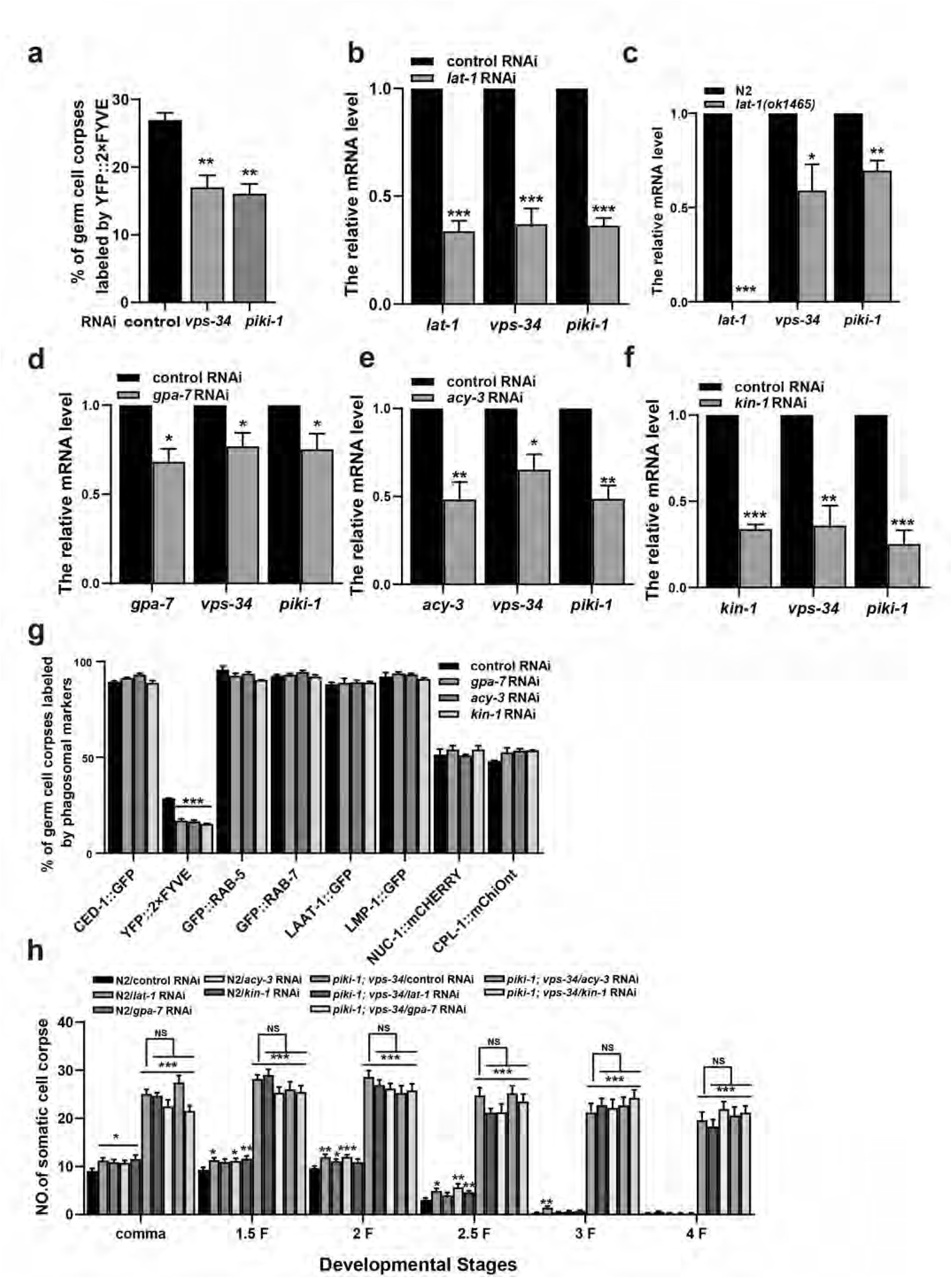
The LAT-1/GPA-7/ACY-3/KIN-1 pathway activates the transcription of VPS-34 and PIKI-1 for phagosomal PI3P generation. (a) Quantification (mean ± SEM) of cell corpse labeling by the phagosomal marker YFP::2×FYVE. A total of ≥100 cell corpses were analyzed, and the data were derived from three independent replicates. Comparisons were performed between the control and *vps-34* or *piki-1* RNAi worms using the YFP::2×FYVE marker. (b–f) The relative mRNA levels of *vps-34* and *piki-1* in the indicated strains were determined by qRT‒PCR. (g) Quantification (mean ± SEM) of the labeling of cell corpses with different phagosomal markers. A total of ≥100 cell corpses were analyzed for each marker, and the data were derived from three replicates. Comparisons were performed among the control, *gpa-7*, *acy-3*, and *kin-1* RNAi worms for each marker. (h) Quantification (mean ± SEM) of embryonic cell corpses at different embryonic stages using the indicated strains. Fifteen embryos were scored at each embryonic stage for each strain. Unpaired t tests were performed. *P < 0.05, **P < 0.01, ***P < 0.001; NS, not significant. All error bars indicate the mean ± SEM.

Because VPS-34 and PIKI-1 act together to drive apoptotic cell degradation, we asked whether they function within the LAT-1/Gs/adenylyl cyclase/PKA pathway. In *piki-1; vps-34* double mutants, additional RNAi targeting *lat-1*, *gpa-7*, *acy-3*, or *kin-1* did not further exacerbate embryonic cell corpse engulfment defects at any stage (Fig. 3h), indicating that *piki-1* and *vps-34* function in the same genetic pathway as *lat-1*, *gpa-7*, *acy-3*, and *kin-1* to promote corpse removal. Together, these findings demonstrate that LAT-1/Gs/adenylyl cyclase/PKA signaling activates the transcription of VPS-34 and PIKI-1 to regulate apoptotic cell clearance.

### AST-1 acts as a transcriptional regulator downstream of LAT-1 to control *vps-34* and *piki-1* expression

Activation of PKA downstream of GPCRs releases catalytic subunits upon cAMP binding, enabling (i) phosphorylation of cytoplasmic targets and (ii) nuclear translocation to phosphorylate CREB, which recruits CBP/p300 to regulate genes bearing cAMP-response elements (CREs)^39^. To identify transcriptional regulators downstream of the LAT-1/Gs/adenylyl cyclase/PKA pathway that activate *vps-34* and *piki-1*, we first assessed CRH-1 (a *C. elegans* CREB homolog) and CBP-1 (CBP/p300 homolog). In *cbp-1* RNAi worms, the number of germline corpses was indistinguishable from that in the controls across three replicates (Table EV8), suggesting that this gene does not play a role in apoptosis. In contrast, *crh-1* inactivation increased the number of germline corpses (Table EV8). When the *xwhIs49 (P_ced-1_rde-1; rde-1)* and *xwhIs82 (P_egl-1_rde-1; rde-1)* strains were used, *crh-1* RNAi significantly increased the number of embryonic corpses in *P_egl-1_rde-1; rde-1* worms but not in *P_ced-1_rde-1; rde-1* worms (Table EV9–10), indicating that *crh-1* primarily affects apoptosis induction rather than corpse removal.

Understanding the transcriptional regulation of these key kinases led us to investigate the downstream transcription factors involved. We therefore performed a secondary RNAi screen of 660 transcription regulator genes to identify potential transcriptional regulators downstream of the LAT-1 pathway (WormBase: transcription regulator activity, GO: 0140110; Table EV11). Inactivation of 72 genes increased the number of germline corpses in at least two replicates (Table EV12). Secondary assays in the *P_ced-1_rde-1; rde-1* and *P_egl-1_rde-1; rde-1* backgrounds revealed 11 genes whose knockdown selectively increased embryonic corpses in *P_ced-1_rde-1; rde-1* worms but not in *P_egl-1_rde-1; rde-1* worms (Table EV13–14), indicating roles in corpse clearance. We hypothesized that the relevant transcriptional regulators would affect the proportion of corpses labeled by YFP::2xFYVE. Indeed, RNAi of *ast-1* or *nhr-168* significantly decreased the percentage of YFP::2xFYVE-positive germline corpses (Fig. 4a). qPCR confirmed that *vps-34* and *piki-1* mRNA expression decreased upon *ast-1* or *nhr-168* knockdown (Fig. 4b–c).

**Fig. 4.**
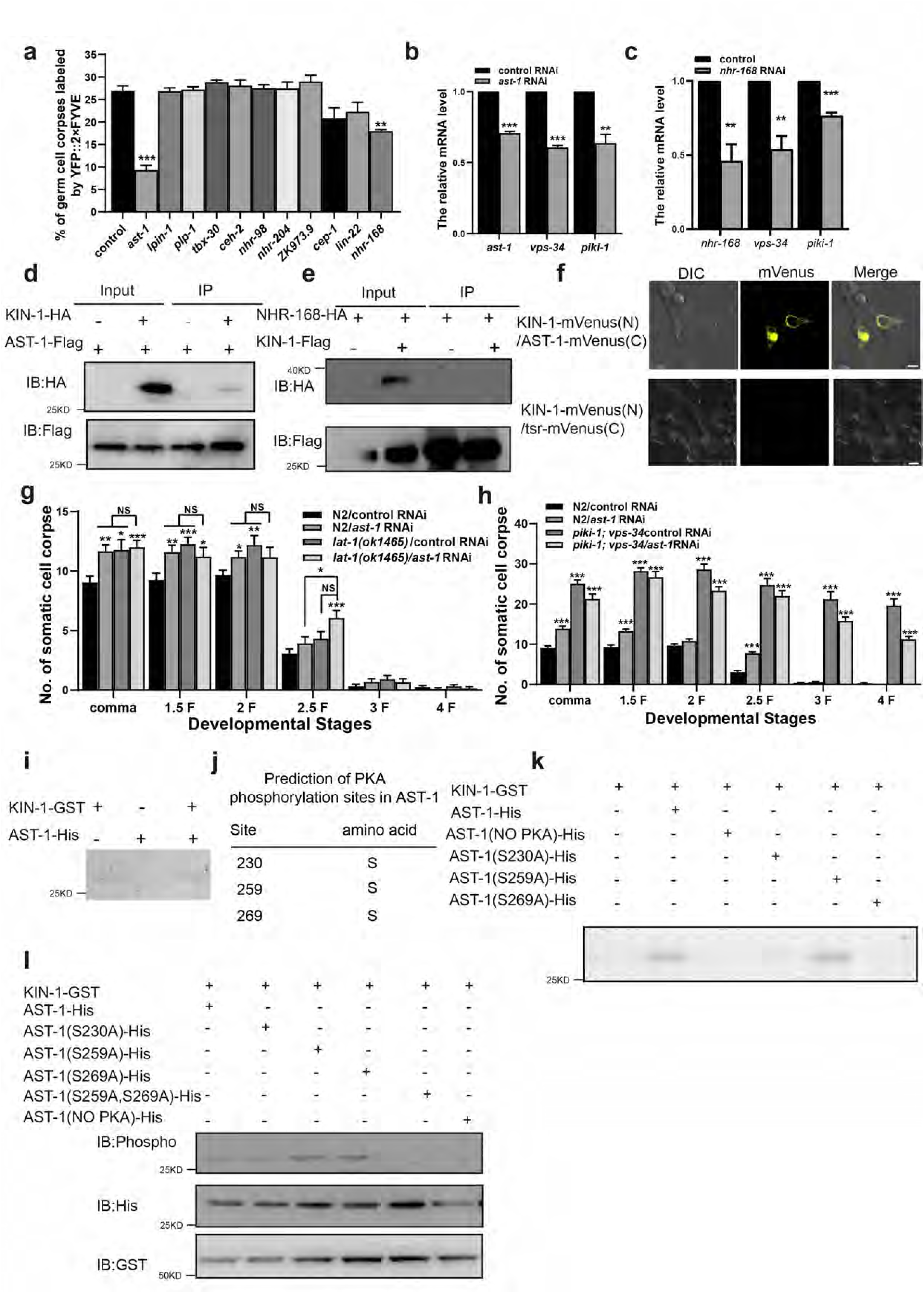
The transcription factor AST-1 functions downstream of LAT-1 to activate the expression of *vps-34* and *piki-1*. (a) Quantification (mean ± SEM) of cell corpse labeling by the phagosomal marker YFP::2×FYVE. A total of ≥100 cell corpses were analyzed, and the data were derived from three replicates. Comparisons were performed between the control and transcription factor RNAi worms using the YFP::2×FYVE marker. (b–c) Relative mRNA levels of *vps-34* and *piki-1* in wild-type (N2) versus (b) *ast-1*(RNAi) or (c) *nhr-168*(RNAi) worms, as analyzed by qRT‒PCR. (d) Co-IP assays in 293T cells to examine the interaction between AST-1 (tagged with FLAG) and KIN-1 (tagged with HA). Proteins were immunoprecipitated using anti-Flag magnetic beads (MBL). (e) Co-IP assays in 293T cells to assess the interaction between KIN-1 (Flag-tagged) and NHR-168 (tagged with HA). The proteins were immunoprecipitated using anti-Flag magnetic beads. (f) Fluorescence images of Venus reconstitution in 293T cells coexpressing KIN-1-mVenus (N) and AST-1-mVenus (C) (top panel). Fluorescence images of 293T cells coexpressing β-actin-mVenus (N) and tsr-mVenus (C) were used as negative controls (bottom panel). Scale bar: 10 μm. (g–h) Quantification (mean ± SEM) of embryonic cell corpses at different developmental stages in the indicated strains. Fifteen embryos were scored at each stage. (i) In vitro kinase assay to detect PKA (KIN-1)-mediated phosphorylation of AST-1. KIN-1 was expressed as a GST fusion protein and purified by GST affinity chromatography. AST-1 was expressed with a His tag and purified via nickel-affinity chromatography. The kinase reaction was performed by incubating purified AST-1 with KIN-1-GST. Following the reaction, AST-1 phosphorylation was detected using Pro-Q Diamond phosphoprotein staining. (j) Schematic representation of the predicted serine PKA phosphorylation sites on AST-1. (k–l) An in vitro kinase assay revealed AST-1 phosphorylation by PKA (KIN-1). WT AST-1-His and the indicated serine-to-alanine mutants were incubated with GST-KIN-1. The reaction products were analyzed by SDS‒PAGE and probed with Pro-Q Diamond phosphoprotein stain (k) or immunoblotted with anti-phospho antibodies (l).

To test whether AST-1 is a direct PKA target, we assessed its interactions with KIN-1. Co-IP demonstrated an interaction between AST-1 and KIN-1 (Fig. 4d), whereas NHR-168 did not interact with KIN-1 (Fig. 4e). While NHR-168 also affected *vps-34* and *piki-1* transcription, its lack of interaction with KIN-1, combined with our focus on the LAT-1/PKA axis, led us to prioritize AST-1 for further investigation in this study. Bimolecular fluorescence complementation (BiFC) in HEK293T cells supported a likely direct interaction between AST-1 and KIN-1 (Fig. 4f). Functionally, *ast-1* RNAi did not further increase embryonic corpse engulfment defects in *lat-1(ok1465)* or in *piki-1;vps-34* double mutants (Fig. 4g–h), placing *ast-1*, *lat-1*, *piki-1*, and *vps-34* in the same genetic pathway to promote corpse removal.

To test whether KIN-1 mediated AST-1 phosphorylation, we performed in vitro kinase assays by incubating AST-1::6XHIS with GST::KIN-1 in a reaction mixture containing ATP. Using Pro-Q Diamond phosphoprotein Gel Stain, we detected the phosphorylation signal when AST-1::6XHIS was incubated with KIN-1 but not in control samples (Fig 4i). To identify the specific phosphorylation sites of AST-1 mediated by KIN-1, we performed in silico prediction using NetPhos 3.1 on the AST-1 protein sequence. Analysis revealed three potential serine phosphorylation sites (S230, S259, and S269) within the conserved motif (residues 230–269) of AST-1 (Fig 4j). To determine the specific phosphorylation sites of AST-1 that are mediated by KIN-1, we generated serine-to-alanine point mutants. In vitro kinase assays revealed that phosphorylation levels, as indicated by both Pro-Q Diamond phosphoprotein gel staining and anti-phosphoserine/threonine antibody detection, were significantly reduced when AST-1 single mutants (S259A or S269A) and double mutants (S259A/S269A) were coincubated with KIN-1. In contrast, this attenuation was not observed in the S230A mutant (Fig 4k-l). To test the effects of abolishing these sites in vivo, we generated transgenic worms with a construct driving the expression of wild-type AST-1, AST-1 single mutants (S259A or S269A), and double mutants (S259A/S269A) in engulfing cells under the control of the CED-1 promoter and subjected them to *ast-1*(5’UTR) RNAi. We found that wild-type AST-1 rescued the engulfment defects observed in *ast-1*(5’UTR) RNAi worms, whereas AST-1 single mutants (S259A or S269A) and double mutants (S259A/S269A) did not rescue the engulfment defects observed in *ast-1*(5’UTR) RNAi worms (Table EV15). These results establish S259 and S269 as the primary phosphorylation sites for KIN-1-mediated phosphorylation of AST-1, which is required for apoptotic cell clearance.

To determine whether AST-1 directly regulates *vps-34* and *piki-1*, yeast one-hybrid (Y1H) assays were used to assess AST-1 binding to their promoters (Fig. EV2a-b). AST-1-AD interacted with *vps-34* promoter-3 (−525 to −281 bp) and *piki-1* promoter-3 (−439 to −174 bp) (Fig. EV2c-d). Sequence analysis revealed four consensus AST-1 sites within the *vps-34* region [(G/A/T)(C/G/A)GGA(A/T)(A/G)] and three within the *piki-1* region. In luciferase reporter assays, AST-1 overexpression increased *piki-1* WT promoter activity 11.2-fold (*p* < 0.001), whereas a site-directed mutation at −269 to −279 bp reduced activity by 90% (*p* < 0.001 vs control); mutations at −332 to −342 bp or −417 to −427 bp did not affect activity (*p* > 0.05). AST-1 similarly increased *vps-34* WT promoter activity 13-fold (*p* < 0.001), while the −315 to −325 bp mutant showed an 80% reduction (*p* < 0.001); mutations at −263 to −273 bp, −279 to −289 bp, or −438 to −448 bp had no significant effect (*p* > 0.05) (Fig. EV2e-f). Electrophoretic mobility shift assays (EMSAs) confirmed direct, sequence-specific binding: purified AST-1::6×HIS formed complexes with biotin-labeled probes spanning the *vps-34* (−305 to −345 bp) and *piki-1* (−249 to −289 bp) promoters. The complexes displayed supershifts with the anti-His antibody and were abolished by excess unlabeled WT—but not mutant—probe (Fig. EV2g-h). Collectively, these data demonstrate that AST-1 binds with high specificity to the −315 to −325 bp region of the *vps-34* promoter and the −269 to −279 bp region of the *piki-1* promoter to drive transcription.

### Engulfment receptor CED-1 physically interacts with LAT-1 to activate downstream gene expression

We next examined how LAT-1 is activated to drive corpse clearance. In vertebrates, teneurins help form transsynaptic complexes by interacting with latrophilins and dystroglycan^40^. *C. elegans* encodes a single teneurin homolog, *ten-1*^41^. *ten-1* RNAi increased the number of embryonic and germline corpses (Fig. EV3a-b). In both the *xwhIs49 (P_ced-1_rde-1; rde-1)* and the *xwhIs82 (P_egl-1_rde-1; rde-1)* backgrounds, *ten-1* RNAi significantly increased the number of embryonic corpses (three replicates; Fig. EV3c-d), suggesting effects on both corpse removal and apoptosis induction. Four-dimensional microscopy revealed prolonged persistence of germline corpses (>100 min) upon *ten-1* knockdown (Fig. EV3e), confirming a clearance defect. Notably, the application of *ten-1* RNAi in engulfing cells exacerbated the defects in embryonic corpse engulfment observed in worms subjected to *lat-1* RNAi treatment in these cells (Fig. EV3f), indicating that *ten-1* does not act in the same genetic pathway as *lat-1* for corpse removal. Furthermore, our Co-IP and BIFC assays did not detect TEN-1–LAT-1 interactions (Fig. EV3h-i), and *ten-1* RNAi did not reduce *vps-34* or *piki-1* transcripts, as determined by qPCR (Fig. EV3g). Together, these data argue against TEN-1 as the activating ligand for LAT-1 in corpse clearance, at least through the specific transcriptional pathway identified here, suggesting that while TEN-1 affects efferocytosis, its mechanism differs from that of LAT-1.

Because *lat-1* functions in the *ced-1/6/7* pathway and TTR-52 connects exposed PS to the CED-1 extracellular domain (ECD), we investigated whether LAT-1 directly recognizes PS. A membrane lipid strip assay did not support PS binding by the LAT-1 ECD (Fig. 5a). Co-IP confirmed the known TTR-52–CED-1 interaction (Fig. 5b) but detected no LAT-1–TTR-52 interaction (Fig. 5c), suggesting that LAT-1 activation requires other receptor(s) or cofactor(s).

**Fig. 5.**
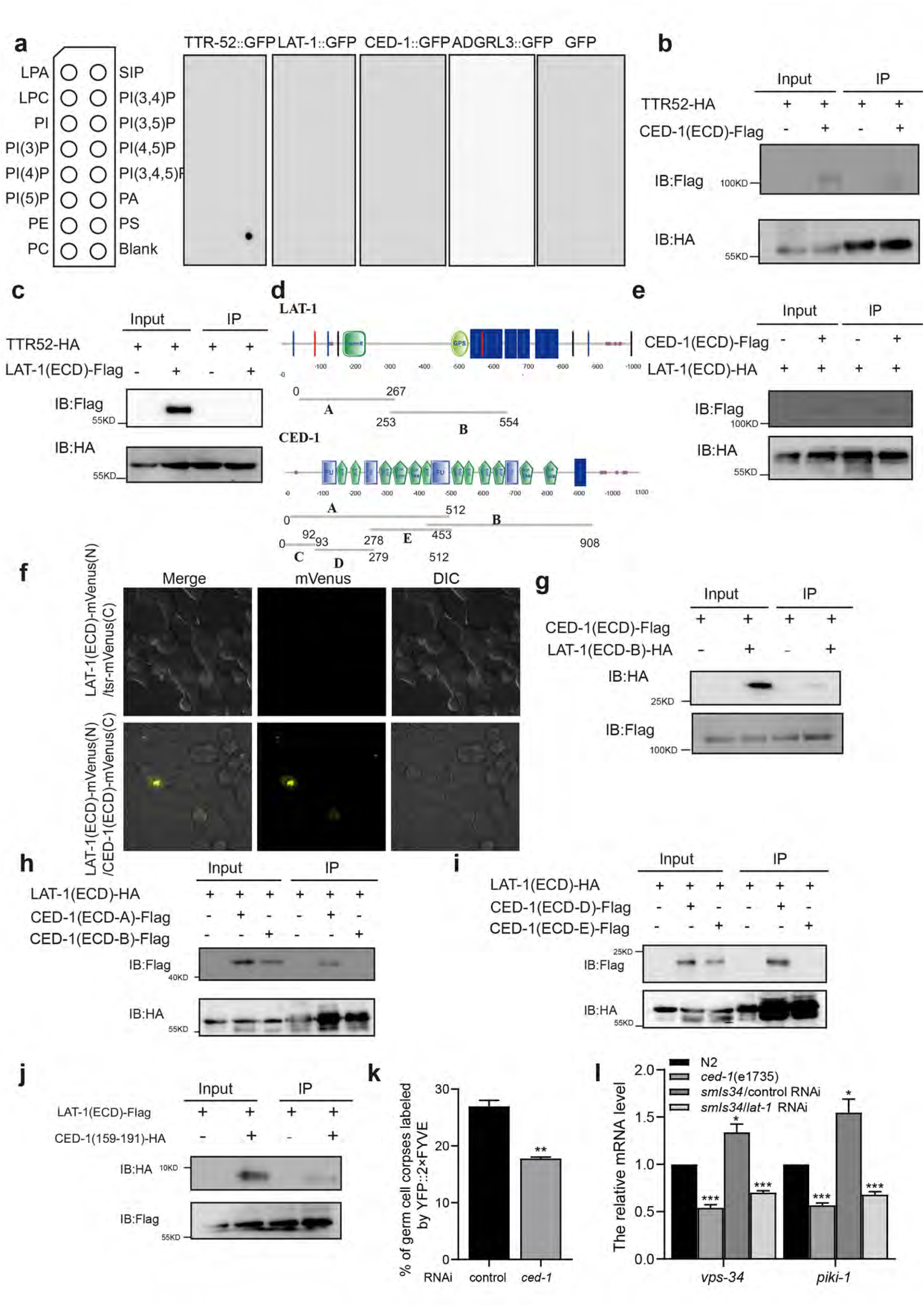
CED-1 directly interacts with LAT-1 to activate the expression of *vps-34* and *piki-1*. (a) Membrane lipid strip assay assessing the binding of the purified LAT-1::GFP extracellular domain to phosphatidylserine (PS) and other indicated lipids. (b) Co-IP assays in 293T cells to assess the interaction between CED-1 (ECD) (Flag-tagged) and TTR-52 (tagged with HA). The proteins were immunoprecipitated using anti-Flag magnetic beads. (c) Co-IP assays in 293T cells to assess the interaction between LAT-1 (ECD) and TTR-52. The proteins were immunoprecipitated using anti-Flag magnetic beads. (d) Schematic representation of LAT-1 and CED-1 protein structures and their respective truncation constructs. (e) Co-IP assays in 293T cells to assess the interaction between CED-1 (ECD) (Flag-tagged) and LAT-1 (tagged with HA). The proteins were immunoprecipitated using anti-Flag magnetic beads. (f) BiFC assay detecting the interaction between the extracellular domains of LAT-1 and CED-1. Scale bar, 10 μm. (g) Co-IP assay testing the interaction between LAT-1 fragment A and the extracellular domain of CED-1. (h) Co-IP assay assessing the interaction between the extracellular domain of LAT-1 and CED-1 fragment B. (i) Co-IP assay evaluating the interaction between the extracellular domain of LAT-1 and CED-1 fragment D. (j) Co-IP assay determining the interaction between the extracellular domain of LAT-1 and CED-1 (amino acids 159–191). (k) Quantification (mean ± SEM) of cell corpse labeling by the phagosomal marker YFP::2×FYVE. A total of ≥100 cell corpses were analyzed, and the data were derived from three independent replicates. (l) Relative mRNA levels of *vps-34* and *piki-1* in wild-type (N2), *ced-1(e1735)*, *smIs34* (which overexpressed a full-length CED-1::GFP fusion protein under its native promoter), and *smIs34*/*lat-1* RNAi worms, as determined by qRT‒PCR. Unpaired t tests were performed. *P < 0.05, **P < 0.01, ***P < 0.001; NS, not significant. All error bars indicate the mean ± SEM.

CED-1 is a single-pass engulfment receptor with multiple extracellular EGF repeats for corpse recognition and an intracellular domain for signaling^17^. LAT-1 contains an RBL domain, an HRM domain, a GPS motif, a 7TM region, and an intracellular domain (ICD)^42^ (Fig. 5d). Co-IP revealed that the LAT-1 ECD interacts with the CED-1 ECD (Fig. 5e). BiFC in HEK293T cells further indicated a direct ECD–ECD interaction (Fig. 5f). Mapping with truncation constructs revealed that CED-1 residues 159–191 are sufficient for binding the LAT-1 ECD and that the LAT-1 GPS motif is required for interaction with the CED-1 ECD (Fig. 5g–j). Additionally, knockdown of *ced-1* led to a substantial decrease in the ratio of YFP::2×FYVE on phagosomes (Fig. 5k). Consistent with a functional link, *ced-1(e1735)* mutants presented significantly reduced *vps-34* and *piki-1* mRNA levels (Fig. 5l). Overexpression of CED-1 in the integrated strain *smIs34* (CED-1::GFP under the native promoter) increased *vps-34* and *piki-1* transcription (Fig. 5l). *smIs34* worms treated with *lat-1* RNAi presented significantly reduced *vps-34* and *piki-1* mRNA levels (Fig. 5l). These observations suggest that the transcriptional activity of *vps-34* and *piki-1* is enhanced by CED-1 overexpression, a process dependent on LAT-1 function.

Our findings support a model in which the CED-1 ECD acts as an activating partner for LAT-1 to activate *vps-34* and *piki-1* transcription, thereby promoting corpse degradation. We subsequently utilized the transgenic integrated strain *smIs110* (CED-1ΔC::GFP, a GFP-fused CED-1 with the C-terminal region deleted) to evaluate this model. The expression of the CED-1 ECD in the integrated strain *smIs110* resulted in decreased transcription levels of *vps-34* and *piki-1*. To eliminate the potential dominant negative effect of CED-1 ECD overexpression on endogenous CED-1, we crossed *smIs110* with *ced-1(e1735)* null allele mutants. Nevertheless, the overexpression of CED-1 ECD did not restore the transcription levels of *vps-34* and *piki-1* in *ced-1(e1735)* worms to those observed in wild-type worms (Fig. EV4a). These findings suggest that the intracellular domain (ICD) of CED-1 is essential for its full function in regulating *vps-34* and *piki-1* transcription, likely by coordinating with downstream effectors in the overall efferocytosis pathway. CED-1 functions as a receptor that binds to apoptotic targets and utilizes its NPXY motif to recruit the adaptor protein CED-6. CED-6 subsequently forms a complex with both clathrin heavy chain (CHC-1) and the AP2 complex (APA-2), which likely facilitates the rearrangement of the actin cytoskeleton necessary for the engulfment of cell corpses. Furthermore, by forming a complex with them, CHC-1 and APA-2 enhance the phagosomal association of LST-4/Snx9/18/33 and DYN-1^43^, thereby promoting phagosome maturation, which is essential for corpse degradation. We detected decreased transcription levels of *vps-34* and *piki-1* in *ced-6(n1813)* mutants as well as in *apa-2*, *chc-1*, and *dyn-1* RNAi-treated worms (Fig. EV4b-e). These findings further indicate that the intracellular domain of CED-1 is necessary for the regulation of *vps-34* and *piki-1* transcription. We next investigated the upstream signals of CED-1 that regulate *vps-34* and *piki-1* transcription during apoptotic cell clearance. The secreted protein TTR-52 dimerizes and binds to phosphatidylserine (PtdSer) exposed on the cell surface. It engages with the extracellular ERM domain of the CED-1 receptor, serving as a molecular bridge that links the “eat me” signal to the CED-1 receptor. We detected decreased transcription levels of *vps-34*, but not *piki-1*, in *ttr-52* RNAi-treated worms and *ttr-52(tm2078)* mutants (Fig. EV4f-g). These findings suggest that the PtdSer “eat me” signal transduced by TTR-52 to CED-1 is required for the transcription of *vps-34*.

### Evolutionary conservation of the CED-1/MEGF10–LAT-1/ADGRL3 regulatory axis in efferocytosis

*C. elegans* LAT-1 is orthologous to *Drosophila* CIRL (calcium-independent receptor for α-latrotoxin) and mammalian latrophilins^44^, whereas CED-1 is orthologous to *Drosophila* Draper (DRPR)^45^. In HEK293T cells, Co-IP of coexpressed CIRL::V5 and DRPR::HA demonstrated an interaction (Fig. EV5a), and BiFC confirmed direct CIRL-DRPR association (Fig. EV5b). When we knocked down *cirl* in apoptotic S2 cells, S2 phagocytes still engulfed several ACs, whereas *xkr*-RNAi ACs (which inhibit PS exposure during apoptosis) significantly reduced the number of engulfed ACs in each phagocyte, indicating that CIRL does not function in efferocytosis in apoptotic cells. Furthermore, we introduced a S2 cell line that expresses caspase-activated GFP and pH-insensitive mCherry (CharON)^46^ to trace apoptosis and efferocytosis. We added apoptotic CharOn cells to control S2 cells and *cirl*-RNAi cells, and the GFP fluorescence signal disappeared gradually as the ACs were degraded by lysosomes, while green and red fluorescence signals were present in most *cirl*-RNAi cells (Fig. EV5c-g). Therefore, *cirl* is required for apoptotic cell degradation in *Drosophila* S2 cells.

Although the results of sequence studies suggest that LAT-1 is most similar to ADGRL1, our phylogenetic analysis results support its orthologous relationship with mammalian ADGRL3 (Fig. EV6a). In RAW264.7 macrophages, *adgrl1* knockdown (as validated by qPCR) did not impair the degradation of apoptotic cells (Fig. EV6b) and did not alter *pik3c3* (*vps-34*) or *pik3c2a* (*piki-1*) mRNA levels (Fig. EV6c). In contrast, *adgrl3* knockdown significantly impaired apoptotic cell degradation in RAW264.7 macrophages (Fig. EV7a-b). To investigate the specific effect of ADGRL3 deficiency on the transcriptional regulation of VPS-34 complex subunits, we analyzed the mRNA expression levels of various complex components following *adgrl3* knockdown (Fig. EV7c). Notably, *adgrl3* shRNA treatment significantly downregulated the expression of PIK3C3 (VPS-34), the core catalytic subunit (*\*\*p < 0.01*). In contrast, the expression levels of other shared core subunits (BECN1 and PIK3R3) and specific markers for distinct complexes (ATG14 for Complex I and UVRAG for Complex II) remained unaffected. These findings reveal a highly specific regulatory mechanism whereby ADGRL3 primarily targets the transcriptional level of PIK3C3 rather than globally altering the expression of the entire VPS-34 complex assembly. To further dissect the signaling cascade linking ADGRL3 to this specific PIK3C3 regulation, we measured intracellular cAMP levels, a canonical downstream effector of adhesion GPCRs, including ADGRL3 (Fig. EV7d). Consistent with the transcriptional data, adgrl3 knockdown significantly reduced cellular cAMP concentrations compared to the control group (\*\*\**p* < 0.001, Fig. EV7d). Re-expression of full-length ADGRL3 in adgrl3-deficient cells fully rescued cAMP production, restoring the levels to those observed in control cells (p > 0.05, not significant). In contrast, the expression of the ADGRL3 ΔGPS mutant, which lacks the G protein-coupling proteolytic site essential for initiating canonical GPCR signaling, failed to rescue the cAMP reduction (\*\*\**p* < 0.001). Together, these results demonstrate that ADGRL3 modulates intracellular cAMP levels in a GPS domain-dependent manner and strongly suggest that this signaling pathway may underlie the specific transcriptional control of PIK3C3, thereby governing VPS-34 complex function without disrupting the broader assembly of the complex.

To further validate the effects of *adgrl1* and *adgrl3* on efferocytosis, apoptotic cells were labeled with TAMRA and cocultured with macrophages^47^. Efferocytosis efficiency was assessed using flow cytometry. The results revealed that knockdown of *adgrl3* impaired the degradation of apoptotic cells, whereas knockdown of *adgrl1* had no significant effect (Fig. EV6d). This finding indicates a specific role for ADGRL3, not ADGRL1, in this conserved pathway. Thus, ADGRL3—not ADGRL1—functions as the mammalian homolog that regulates apoptotic cell clearance, potentially via transcriptional control of phosphoinositide kinases.

To further validate the physiological role of ADGRL3, we generated *adgrl3* global knockout mice via gene targeting (Fig. EV8a). We introduced a frameshift mutation into the ADGRL3 coding region, resulting in premature translation termination. Gene expression and protein levels were analyzed in wild-type (*adgrl3^+/+^*), heterozygous (*adgrl3^+/-^*), and homozygous knockout (*adgrl3^-/-^*) mice. Quantitative RT‒PCR analysis revealed a significant ∼60% reduction in *adgrl3* mRNA levels in *adgrl3^-/-^* mice compared with those in control mice (*p* < 0.001; Fig. EV8b). Consistent with these findings, western blot analysis (Fig. EV8c) confirmed that ADGRL3 protein was nearly absent in *adgrl3^-/-^* mice. Notably, the loss of ADGRL3 was accompanied by a marked downregulation of both PIK3C3 and PIK3C2A protein levels. Additionally, we observed decreased protein levels of PIK3C3 and PIK3C2A in *adgrl3*^⁻/⁻^ mice (Fig. EV8c-d). Bone marrow–derived macrophages (BMDMs) from these mice were incubated with UV-induced apoptotic Jurkat cells expressing the genetically encoded fluorescent reporter CharON for 2 h (Fig. EV8e-f), followed by time-course confocal analysis (Fig. EV8f). Loss of ADGRL3 did not affect apoptotic cell uptake but significantly impaired corpse degradation in *adgrl3*^⁻/⁻^ BMDMs (Fig. 6a–c).

**Fig. 6.**
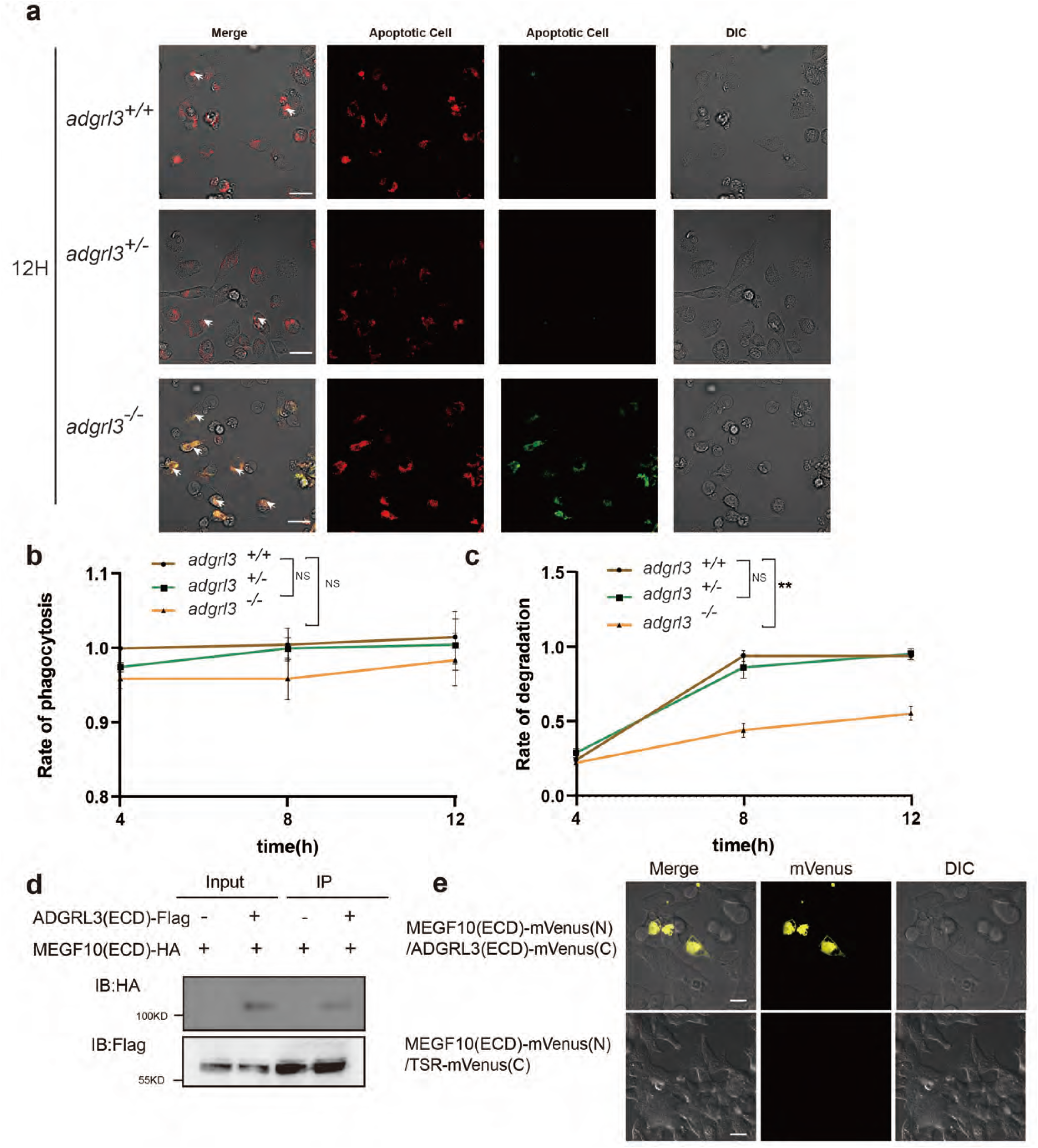
The function and regulatory mechanism of LAT-1 in efferocytosis are evolutionarily conserved in mammals. (a) Effects of *adgrl3* knockout on the degradation of apoptotic cells in BMDMs. The data are representative of at least three independent experiments. For each experiment, five random imaging fields were analyzed. *P < 0.05, **P < 0.01, ***P < 0.001; NS, not significant. Scale bars, 10 μm. (b–c) Time course of apoptotic cell clearance. Phagocytosis (b) and degradation (c) rates were quantified in control (*adgrl3^+/+^*), ADGRL3-heterozygous (*adgrl3^+/—^*), and ADGRL3-KO (*adgrl3^—/—^*) BMDMs at 4, 8, and 12 hours. (d) Co-IP assays in 293T cells to assess the interaction between ADGRL3 (ECD) and MEGF10. Proteins were immunoprecipitated using anti-HA-magnetic beads. (e) BiFC assay detecting the interaction between the extracellular domains of ADGRL3 and MEGF10 in 293T cells. Scale bar, 10 μm.

Finally, to investigate the evolutionary conservation of the LAT-1–CED-1 regulatory module, we examined the interactions between the mammalian homologs of these proteins. We detected a lack of binding between the extracellular domains of ADGRL3 and PS (Fig. 5a), which is consistent with the results of the PS binding experiments with LAT-1, the homolog of ADGRL3 in nematodes. To elucidate the specific structural region of MEGF10 responsible for its interaction with ADGRL3, we first performed three-dimensional structural modeling to predict potential interaction interfaces. Through this modeling, we constructed a spatial configuration of the MEGF10 (green) and ADGRL3 (magenta) complexes (Fig. EV9a, left panel). The overall model revealed that MEGF10 adopts an elongated claw-like structure, whereas a specific domain of ADGRL3 forms a tight binding interface with it. Further high-magnification analysis (Fig. EV9a) revealed the microscopic contact interface: in the region closest to the interaction site, several key amino acid residues of MEGF10 (N168, T182, D165, A170, G154, K162, and K150) and corresponding residues on ADGRL3 (A444, R454, K511, K513, K516, and L517) engage in dense spatial complementarity and molecular contacts. This high-resolution docking simulation visually localizes the critical physical determinants driving their association. We subsequently mapped these three-dimensional structural predictions onto the linear domain architecture diagrams of the respective proteins (Fig. EV9b). Structural analysis of MEGF10 indicated that the region involved in these tight contacts is highly concentrated within its multiple EGF-like domains (depicted by green arrowheads). These structural predictions suggest that specific EGF-like domains (s) of MEGF10 may constitute the core scaffold required for binding to ADGRL3. Co-IP assays in HEK293T cells coexpressing ADGRL3::FLAG and MEGF10::HA (the mammalian ortholog of CED-1) confirmed a physical interaction between the proteins (Fig. 6d), which was further confirmed by BiFC, indicating a direct ADGRL3–MEGF10 association (Fig. 6e). A series of truncation mutants of MEGF10 were subsequently generated, and their binding capacity to ADGRL3 was evaluated, revealing that the second EGF-like domain of MEGF10 is critical for this interaction (Fig. EV9c-e). Functional studies in RAW264.7 macrophages demonstrated that the overexpression of ADGRL3, combined with the exogenous addition of a peptide corresponding to the EGF domain of MEGF10, significantly increased the clearance of apoptotic cells (Fig. EV9f).

To further determine the functional role of *megf10* in phagocytic degradation, we performed knockdown experiments using RAW264.7 cells. Depletion of MEGF10 impaired the degradation of apoptotic cells (Fig. EV10a-b) and concomitantly reduced the expression levels of *pik3c3* and *pik3c2a* at both the transcriptional and protein levels (Fig. EV10c-f). Collectively, these data demonstrate that both the function of LAT-1/ADGRL3 in efferocytosis and its regulatory mechanism via a CED-1/MEGF10-like receptor to control VPS-34/PIKI-1 (and mammalian homologs) are conserved from worms to flies to mammals.

To identify the specific mammalian ortholog of AST-1, we first conducted a phylogenetic analysis, which revealed that *ast-1* is closely related to FLI1 and ETV1 in mammals (Fig. 7a). We then evaluated the functional roles of the two leading candidates, ETV1 and FLI1. Confocal microscopy revealed that knockdown of *etv1*, but not *fli1*, significantly impaired apoptotic cell degradation in RAW264.7 macrophages (Fig. 7c). Consistent with this functional defect, *etv1* knockdown also led to decreased expression of *pik3c3* and *pik3c2a* (Fig. 7b). These results identified ETV1 as a functional ortholog of AST-1 in macrophages. To determine whether the functional conservation of the AST-1 ortholog ETV1 extends to the ADGRL3 pathway, we investigated whether its overexpression could rescue the *adgrl3*^⁻/⁻^ phenotype. Indeed, ETV1 expression in *adgrl3*^⁻/⁻^ BMDMs ameliorated the degradation defect, although not to the level observed in *adgrl3^+/+^* BMDMs (Fig. 7d–e), indicating the evolutionary conservation of function within this signaling axis. To further investigate whether ETV1 participates in apoptotic cell clearance in its phosphorylated form, we performed bioinformatic analyses of its secondary structure and potential phosphorylation sites and identified two conserved serine residues (S334 and S390) as putative regulatory sites for ETV1 activation. To determine the specific phosphorylation sites of PRKACA, the ortholog of *C. elegans* KIN-1, we generated serine-to-alanine point mutants of ETV1. Our in vitro kinase assays revealed that PRKACA directly phosphorylates ETV1 at both Ser334 and Ser390, indicating that these posttranslational modifications are involved in the regulation of ETV1 function during efferocytosis (Fig. EV11a). Notably, both single mutations impaired the nuclear translocation of ETV1, suggesting a role for these phosphorylation events in subcellular localization (Fig. EV11b). Furthermore, in *etv1*-knockdown RAW264.7 cells, reconstitution with wild-type ETV1 restored phagocytic capacity, whereas the expression of the S344A mutant failed to fully rescue the engulfment defect, and the S390A mutation only partially restored phagocytosis (Fig. EV11c). These results establish S334 and S390 as primary phosphorylation sites for PRKACA-mediated phosphorylation of ETV1, which is important for regulating ETV1 function during efferocytosis. Although the amino acid phosphorylation sites of AST-1 by KIN-1 and those of ETV1 by PRKACA are not particularly conserved in their amino acid sequences, our analysis of their three-dimensional protein structures revealed that the phosphorylation sites of AST-1 and ETV1 by PRKACA are highly conserved (Fig. EV12a-b), which demonstrated that the phosphorylation of AST-1/ETV1 by KIN-1/PRKACA to regulate efferocytosis is evolutionarily conserved.

**Fig. 7.**
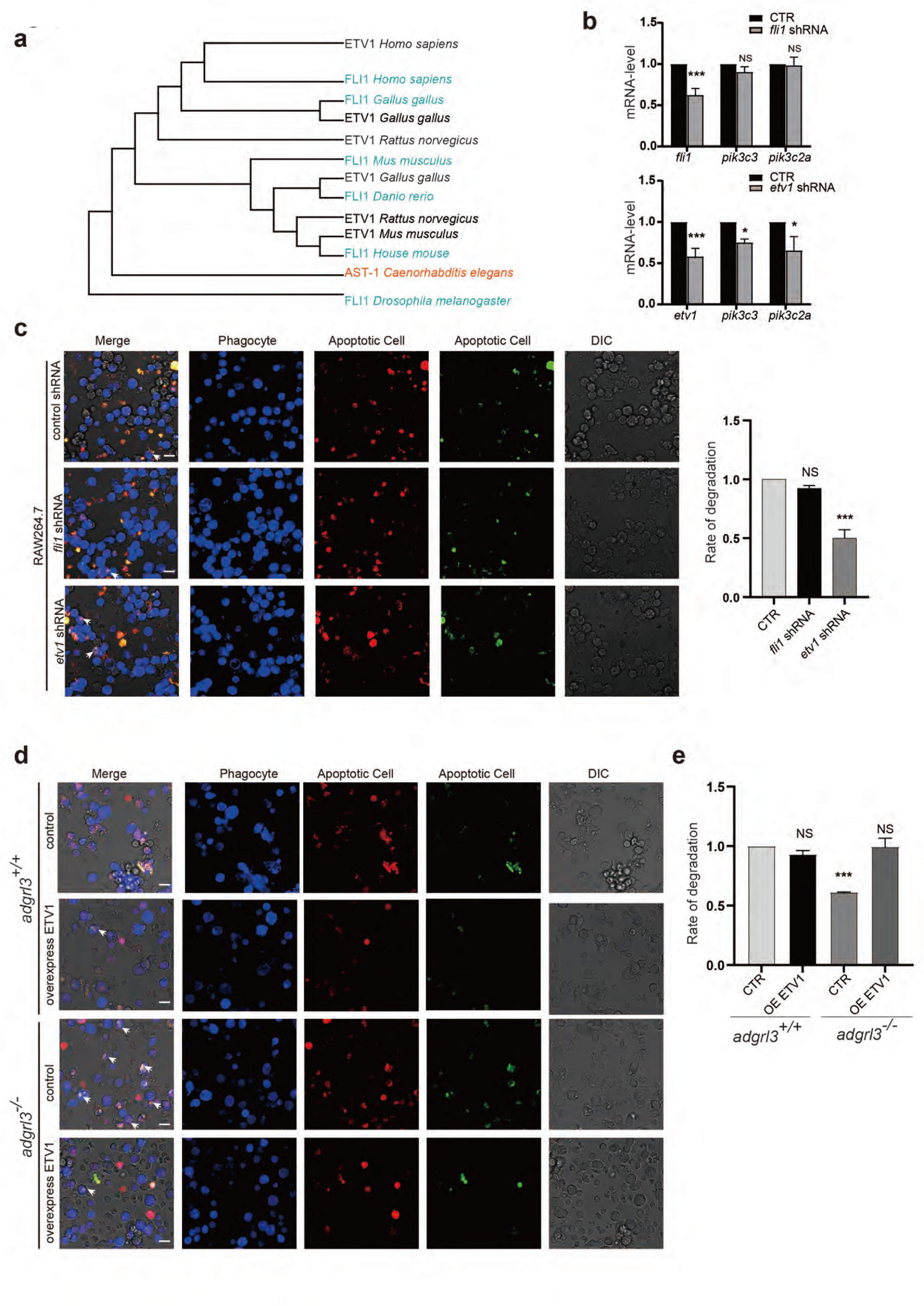
*etv1* is involved in apoptotic cell degradation. (a) Phylogenetic tree of AST-1 orthologs constructed using the maximum likelihood method across representative species. (b) Effect of *etv1* and *fli1* knockdown on the relative mRNA levels of *pik3c3* and *pik3c2a*. The data are representative of at least three independent experiments. *P < 0.05, **P < 0.01, ***P < 0.001; NS, not significant. (c) Roles of *etv1* and *fli1* in macrophage degradation of apoptotic cells. Macrophages expressing control or *etv1*/*fli1*-targeting shRNA (blue) were incubated with dual-color apoptotic CharON Jurkat reporters (green: apoptosis-induced, acid-sensitive; red: acid-resistant). Scale bars, 10 μm. (d–e) Effect of ETV1 overexpression on degradation efficiency in ADGRL3-KO (*adgrl3*^—/—^) BMDMs. Values are mean ± SEM. One-way ANOVA followed by post-hoc test was used. *P < 0.05, **P < 0.01, ***P < 0.001; NS, not significant. Scale bars, 10 μm.

## Discussion

To explore whether GPCRs participate in efferocytosis, we performed a systematic RNAi knockdown of 709 *C. elegans* GPCR genes and identified *srw-40*, *lat-1*, *seb-3*, *sra-11*, and *W08G11.5* as candidates affecting apoptotic cell clearance rather than apoptosis induction. Among these, *lat-1* knockdown caused a pronounced accumulation of cell corpses, prompting further analysis. The functions of *srw-40*, *seb-3*, *sra-11*, and *W08G11.5* in efferocytosis remain to be elucidated and represent important directions for future investigations.

*C. elegans* harbors two latrophilin homologs, *lat-1* and *lat-2*, with *lat-1* more closely related to mammalian latrophilin^30^. Although their functions partially overlap, only *lat-1* is essential for normal development. Consistent with these findings, *lat-2* knockdown did not affect apoptosis, confirming that *lat-1* specifically mediates cell corpse clearance. A recent study suggested that *lat-1* regulates sperm guidance, ovulation, and germ apoptosis non–cell-autonomously, as receptor expression on gonadal sheath cells rescued the increased germline apoptosis phenotype. The authors hypothesized that the increased number of apoptotic germ cells reflected defective corpse clearance rather than elevated apoptosis initiation^48^. Our findings support this hypothesis: *lat-1* activity in engulfing cells was crucial for cell corpse clearance, and *lat-1* knockdown or mutation significantly prolonged corpse persistence without altering the overall apoptosis frequency. Thus, LAT-1 participates in apoptotic cell clearance but not in apoptosis induction.

Several GPCRs are known to regulate efferocytosis. For instance, phosphatidylserine (PS) on apoptotic cells binds to the N-terminal thrombospondin-like repeats of BAI1, triggering ELMO–Dock180–Rac signaling to promote engulfment^27^. GPR132 mediates efferocytosis-induced macrophage proliferation (EIMP) through lactate signaling, which stabilizes Myc^28^. Moreover, macrophage GPR37 senses prosaposin-bearing extracellular vesicles to increase TIM4 expression via the ERK–AP1 axis, thereby increasing efferocytic efficiency^29^. In contrast to these mechanisms, LAT-1—although also an adhesion GPCR—regulates apoptotic cell degradation rather than engulfment. This functional distinction clearly differentiates LAT-1 from other adhesion GPCRs, such as BAI1, which primarily facilitate the initial recognition and internalization of apoptotic cells. Our findings thus reveal a novel role for adhesion GPCRs in the postengulfment stages of efferocytosis.

PtdIns3P accumulation on phagosomes is essential for corpse degradation^37^. We found that *lat-1* disruption reduced the recruitment of the early phagosome marker PtdIns3P, indicating a role in early phagosome maturation. Given that PtdIns3P also governs endocytic trafficking, retrograde transport, and autophagy, determining whether LAT-1 contributes to these additional PtdIns3P-dependent processes is important. Such investigations could reveal broader roles for LAT-1 in membrane trafficking beyond efferocytosis.

Latrophilins, mammalian homologs of *lat-1*, belong to the adhesion GPCR (aGPCR) family, the second largest class of GPCRs^49,50^. Early studies of ADGRL1 (LPHN1) demonstrated that α-latrotoxin binds and couples to Gαo, leading to elevated cAMP and IP₃ levels^50,51^. In *C. elegans*, LAT-1 regulates anterior–posterior cell division alignment via a Gs/adenylyl cyclase–dependent cAMP pathway. Consistent with these precedents, our results demonstrate that LAT-1 mediates apoptotic cell clearance through a canonical GPCR-G protein signaling cascade. Specifically, LAT-1 couples to GPA-7 (a Gα subunit) and adenylyl cyclase, thereby activating protein kinase A (PKA). This signaling architecture establishes LAT-1 as a bona fide GPCR that transduces extracellular signals into intracellular second messenger cascades during efferocytosis.

The PI3 kinases PIKI-1 and VPS-34 sequentially regulate phagosome maturation by orchestrating PtdIns3P dynamics, which is counteracted by the PI 3-phosphatase MTM-1. PIKI-1 and VPS-34 play distinct temporal roles: PIKI-1 acts during phagosome sealing and early PtdIns3P formation, whereas VPS-34 drives later enrichment on sealed phagosomes^38^. We identified AST-1, an ETS-domain transcription factor, as a phosphorylation substrate of KIN-1 that directly regulates *vps-34* and *piki-1* transcription. These findings reveal that LAT-1 controls vps-34 and piki-1 expression through a Gs/adenylyl cyclase/cAMP/PKA/AST-1 signaling cascade, thereby ensuring adequate levels of these critical enzymes for efficient efferocytosis. This transcriptional regulatory mechanism represents a previously unrecognized layer of control in phagosome maturation. Beyond its role in efferocytosis, AST-1 controls axon navigation, differentiation of dopaminergic neurons, and pharyngeal morphogenesis^53,54^. Whether the LAT-1/PKA/AST-1 pathway contributes to these AST-1-dependent developmental processes remains an important question for future investigation. Such studies could reveal whether this signaling axis represents a general mechanism through which LAT-1 regulates diverse biological processes through transcriptional control.

Endogenous membrane-tethered teneurins bind the extracellular regions of ADGRL1–3 and promote intercellular adhesion and synapse formation^50^. The sole *C. elegans* teneurin homolog, *ten-1*, is essential for tissue organization and neural development^55^. We found that *lat-1* and *ten-1* do not function as receptor–ligand partners during corpse clearance, which is consistent with their partially overlapping but distinct developmental roles. A recent study revealed that TOL-1, the sole Toll-like receptor in *C. elegans*, serves as a ligand for LAT-1. The interaction between TOL-1 and LAT-1 is essential for the early developmental stages of *C. elegans*^49^. Another study demonstrated that LAT-1 directly interacts with the DSL protein/Notch ligand LAG-2, thereby enhancing Notch receptor activation to modulate cell proliferation within the gonadal stem cell niche^55^. These findings indicate that LAT-1 engages multiple ligands to participate in diverse biological processes.

In the context of efferocytosis, our structure‒function analyses revealed that LAT-1 contains an RBL domain, an HRM domain, a GAIN (GPCR autoproteolysis-inducing) domain, a seven-transmembrane (7TM) region, and an intracellular domain. We show that a single EGF-like repeat in CED-1 engages the GAIN domain of LAT-1, likely triggering Stachel-mediated receptor activation. Because the GAIN domain functions as an agonistic module necessary for receptor activation, binding of the CED-1 EGF repeat may induce conformational changes that expose the Stachel sequence to initiate LAT-1 signaling. Future structural studies should focus on elucidating how this interaction is triggered at the molecular level and whether specific 7TM residues serve as coactivation sites.

Our findings demonstrate that CED-1 regulates *vps-34* and *piki-1* transcription through LAT-1 activation, linking receptor engagement to transcriptional control (Fig. EV13). Interestingly, the intracellular domain of CED-1 is also crucial for its complete ability to regulate *vps-34* and *piki-1* transcription, likely through coordination with downstream effectors within the broader efferocytosis pathway. To identify upstream signals that regulate CED-1-mediated transcriptional control, we examined the role of TTR-52, which mediates the recognition of phosphatidylserine (PtdSer). We observed reduced *vps-34* transcription in both *ttr-52* RNAi-treated worms and *ttr-52(tm2078)* mutants, whereas *piki-1* levels remained unchanged. These findings imply that the PtdSer “eat-me” signal, conveyed by TTR-52 to CED-1, plays a crucial role in facilitating *vps-34* transcription specifically. Beyond facilitating PtdSer recognition on apoptotic cells, TTR-52 collaborates with CED-7 and NRF-5 to mediate PtdSer transfer from apoptotic cells to adjacent phagocytes, thereby enhancing recognition and engulfment^56^. One possible explanation for the selective effect of TTR-52 on *vps-34* transcription, as opposed to *piki-1*, is that TTR-52 may regulate an as-yet-unidentified signaling molecule within phagocytes that specifically modulates *piki-1* transcriptional regulation. The ABC transporter CED-7, operating within the same genetic pathway as CED-1 and TTR-52, is crucial for the localization of CED-1 around apoptotic cell debris. The precise role of CED-7 in facilitating the recognition of cell corpses by CED-1 remains unclear. Notably, the extracellular domain of CED-1 can bind to phosphatidylserine (PtdSer) in vitro, suggesting that CED-1 may detect cell corpses through direct interaction with PtdSer^57^. Therefore, it is plausible that multiple mechanisms contribute to the recognition process of cell corpses by the CED-1 receptor. Further investigation is needed to elucidate the upstream signals that regulate CED-1-mediated transcriptional control of vps-34 and piki-1 during apoptotic cell clearance.

Previous work has established that the CED-1 pathway governs both engulfment and degradation. During the degradation phase, CED-1 remains on nascent phagosomes and cooperates with CED-6 and DYN-1 to directly recruit PIKI-1 and VPS-34, promoting PtdIns3P production and early phagosome maturation^21^. Our findings extend this model by revealing an additional transcriptional mechanism: CED-1 also activates *vps-34* and *piki-1* expression through LAT-1 signaling to ensure sufficient enzyme levels for efficient degradation (Fig. EV13). This dual regulatory strategy—combining direct protein recruitment with transcriptional upregulation—ensures a sufficient pool of VPS-34 and PIKI-1 for robust phagosome maturation. This concept is consistent with reports that CHN-1, a U-box E3 ubiquitin ligase, catalyzes K63-linked ubiquitination to stabilize VPS-34 protein levels^58^. Whether PIKI-1 is similarly regulated by posttranslational mechanisms remains to be determined. Collectively, our data establish LAT-1 as a specific coreceptor of CED-1 that promotes apoptotic cell degradation by activating *vps-34* and *piki-1* transcription, complementing the direct recruitment mechanism to ensure adequate enzyme availability.

LAT-1 shares homology with *Drosophila* CIRL and mammalian ADGRL3 (LPHN3)^45^. CIRL modulates mechanosensory neuron activity and planar cell polarity via Toll-8^59^, whereas ADGRL3 regulates hippocampal synaptogenesis and has been genetically linked to ADHD^60^. We demonstrate that LAT-1 function in efferocytosis is conserved from nematodes to mammals. CIRL was required for efferocytosis in vitro, and ADGRL3 controlled apoptotic cell degradation in macrophages—providing the first direct mechanistic evidence that CIRL and ADGRL3 function in efferocytosis.

We further showed that CIRL directly interacts with DRPR, the *Drosophila* CED-1 ortholog^60^, MEGF10 and MEGF11, the mammalian CED-1 ortholog^61^. The homologs of *lat-1*, *ced-1*, and *ast-1* also regulate *vps-34* and *piki-1* orthologs in mammalian cells, strongly suggesting that the CED-1–LAT-1 regulatory axis, including its transcriptional mechanism, is evolutionarily conserved. While our data provide robust evidence for key interactions and functional conservation, future in vivo studies are crucial to fully verify the operation of the entire signaling cascade in mammalian organisms.

In addition to MEGF10, JEDI-1 has been directly identified as a homolog of CED-1, sharing conserved domain architecture and signaling mechanisms^62^. MEGF10 facilitates the engulfment of apoptotic neurons by astrocytes and glial precursors^63^. JEDI-1 is crucial for the engulfment of apoptotic sensory neurons during mammalian development ^64^. Future research should determine whether JEDI-1 functions similarly to MEGF10 in activating ADGRL3 for apoptotic cell degradation during mammalian development, which would further elucidate the generality of this regulatory mechanism.

Class III PI3K complexes are centered around the lipid kinase VPS-34 and predominantly exist as two distinct assemblies. VPS-34 complex I comprises VPS-34, VPS15, Beclin1, and ATG14 (ATG14L/Barkor) and is frequently accompanied by NRBF2, which is essential for the initiation of autophagy, including autophagosome nucleation and PI(3)P production on isolation membranes. Conversely, VPS-34 complex II consists of VPS-34, VPS15, Beclin1, and UVRAG (Vps38) and is involved in endosome‒lysosome trafficking, endocytic sorting, and autophagosome maturation. Our findings indicate that ADGRL3 is necessary for the transcriptional regulation of *pik3c3* (encoding VPS-34) and *pik3c2a* (encoding a PIKI-1 ortholog). Importantly, ADGRL3 does not regulate other components of class III PI3K complexes, thereby demonstrating the specific regulatory influence of ADGRL3 on catalytic kinase subunits rather than the entire complex. LC3-associated phagocytosis (LAP) involves a noncanonical autophagy pathway that facilitates the efficient and immunological degradation of apoptotic cells by phagocytes^65^. LAP couples the phagocytic recognition of apoptotic cells (via PtdSer receptors) to rapid, ROS-dependent LC3 lipidation. Rubicon (RUBCN) serves as a pivotal, nonredundant regulator of LAP, recruiting and stabilizing the VPS-34-Beclin1-UVRAG complex on the phagosome to produce sustained PI(3)P, which is essential for LC3 lipidation^66^. An intriguing area for future investigations would be to explore whether ADGRL3-mediated transcriptional regulation of VPS-34 contributes to LAP efficiency, potentially by linking GPCR signaling to this specialized form of phagocytosis.

A recent study revealed that phosphatidylinositol 3,5-bisphosphate facilitates synaptic vesicle cotransport and presynapse assembly^67^. Given that ADGRL3 controls *vps-34* and *piki-1*, two PtdIns 3-kinases essential for generating PtdIns3P, determining whether ADGRL3-mediated transcription of these kinases contributes to synapse formation will be interesting.

In summary, our findings establish that LAT-1 mediates apoptotic cell degradation through a canonical GPCR–G protein signaling cascade that promotes phagosome maturation in *C. elegans*. LAT-1 activates the Gs protein/adenylyl cyclase/PKA/AST-1 pathway, driving *vps-34* and *piki-1* transcription for PtdIns3P production on phagosomes. We further demonstrated that CED-1/MEGF10 directly engages LAT-1/ADGRL3 via its extracellular EGF-like repeat to stimulate this pathway (Fig. EV9c-e). Moreover, a peptide corresponding to the EGF domain of MEGF10 triggered ADGRL3 receptor function for apoptotic cell degradation (Fig. EV9f). The conserved LAT-1/ADGRL3–CED-1/MEGF10 axis represents a central mechanism of efferocytosis across species.

This regulatory pathway has potential clinical relevance: defective efferocytosis contributes to the pathogenesis of autoimmune diseases such as systemic lupus erythematosus (SLE) and neurodegenerative disorders such as Alzheimer’s disease. Thus, modulating LAT-1/ADGRL3–mediated efferocytosis could provide novel therapeutic opportunities for these conditions. Future in vivo studies are crucial to fully elucidate the protective role of the CED-1/MEGF10–LAT-1/ADGRL3 axis against autoimmune and neurodegenerative diseases and to explore its therapeutic potential.

## Materials and Methods

### *C. elegans* strains and genetics

The Bristol strain N2 of *Caenorhabditis elegans* was used as the wild type. Strains of *C. elegans* were cultured at 20°C on Nematode Growth Medium (NGM) and maintained at that temperature using standard protocols. Other mutant alleles used in this study are listed according to the linkage groups: LGI: *ced-1(e1735)*, *ced-2(n1994)*; LGII: *lat-1(ok1465)*; LGIII: *ced-6(n1813)*; LGIV: *ced-3(n717)*, *ced-5(n1812)*. All the mutants were outcrossed at least four times with the N2 strain. Nematode strains registered in our laboratory at WormBase were designated SNU (strain) and *xwh* (allele). Transgenic animals carrying extrachromosomal arrays (*xwhEx*) were generated using standard microinjection methods, and integrated genome arrays (*xwhIs*) were generated by UV irradiation to achieve stable expression from arrays with low copy numbers. All the strains used in this study are listed in Table EV16.

### RNAi experiments

Bacterial feeding assays were used in the RNAi experiments. We selected 8–15 wild-type N2 worms in the egg laying stage and placed them in culture medium containing both control and target gene RNAi for 2–3 hours to induce egg laying, after which the females were removed. The worms were then placed in a 20°C incubator for 48 hours until they reached the L4 stage. The L4-stage worms were then transferred to fresh RNAi medium. After 48 hours, the number of embryos laid and apoptotic germ cells were counted.

### Quantification of Embryonic and Gonadal Cell Corpses

To prepare 2% agarose slides, Levamisole solution (10 µl) was added to the slides. Adult worms or embryos were transferred onto slides and observed under a DIC microscope. For embryo cell corpses, at least 15 embryos at each developmental stage (comma, 1.5-fold, 2-fold, 2.5-fold, 3-fold, and 4-fold) in each strain were scored as head regions of embryonic cell corpses. Cell corpses in the germline meiotic region of one gonad arm in each of at least 15 animals were scored at the indicated adult ages (12, 24, 36, 48, and 60 h after the L4 larval stage). The procedure was repeated three times, and the number of apoptotic cells in the gonads or embryos was counted. Time-lapse Imaging and Microscopy Four-dimensional microscopy was used to capture the persistent presence of apoptotic cells. First, embryos (at the two- or four-cell stage) were isolated from hermaphrodites capable of laying eggs. The eggs were placed in an embryonic saline solution and pipetted onto preprepared agar slides. To maintain the survival environment of the eggs, the coverslips were sealed with beeswax, and the imaging temperature was maintained at 20°C. Apoptotic cells were imaged using a Zeiss optical microscope, and 20–30 Z-axis images were captured every 60 s over a 400 min recording period.

Worms tagged with fluorescence were imaged by DIC and fluorescence using an Axio Imager M2 microscope (ZEISS). The images were processed and viewed using ZEN 2 Pro software (Zeiss). Immersol 518F oil (Zeiss) was used for immersion. All the images were captured at 20°C.

### Quantification of Phagosomal Markers

The transgenic worms carrying phagosomal markers that were treated with control and target gene RNAi were mounted on 2% agar pads. Images of the total number of cell corpses and the number of cell corpses that were labeled with different phagosomal markers were captured using an Axio Imager M2 microscope (Zeiss). The percentage of embryonic or gonadal cell corpses labeled by phagosomal markers was determined by dividing the number of labeled cell corpses by the total number of cell corpses.

### AO Staining

Apoptotic cells in the gonads of *C. elegans* were stained with acridine orange (AO). Adult worms were incubated at room temperature in the dark for 2 hours in M9 medium supplemented with acridine orange (100 μg/ml) and OP50 bacteria. The worms were then transferred to Nematode Growth Medium substrate for a 1-h recovery period to reduce background intestinal fluorescence. We observed stained *C. elegans* under a fluorescence microscope and determined the percentage of positively stained cells. Three independent experiments were performed to ensure the scientific validity of the results.

### Tracking apoptotic cells and the duration of exogenous GFP signals

The transgenic worm *ujIs113*(HIS::mCherry) was used to track the degradation time of apoptotic cells in the gonads of *C. elegans*. First, approximately 15 transgenic worms, *ujIs113,* were placed in control and target gene RNAi for egg laying. After 3 hours, the hermaphrodites were removed and placed in a 20°C incubator for 48 hours to develop to the L4 stage. At this stage, the worms were transferred to fresh medium for continued development for 48 hours. Adult worms were then placed on agarose slides containing levamisole. Using fluorescence microscopy, new apoptotic cells were identified, and the process from initiation to disappearance was captured and observed.

The uptake and degradation of exogenous proteins were studied in the transgenic worm line arIs36 (PhspssGFP). *arIs36* was used to track the uptake and degradation of soluble ssGFP secreted by cavity cells. First, approximately 15 hermaphrodite *C. elegans arIs36* individuals were placed in control and target gene RNAi for egg laying. After 3 hours, the hermaphrodites were removed and placed in a 20°C incubator for 48 hours to develop to the L4 stage. At this stage, the nematodes were transferred to fresh medium for continued development. After 48 hours, the culture plates were subjected to heat shock at 33°C for 1 hour in a water bath, followed by recovery and continued growth at 20°C. At designated time points (1 h, 6 h, 12 h, 24 h, 36 h, and 48 h postrecovery), the ssGFP signals in the body cavity and body cavity cells were observed using fluorescence microscopy, and images were captured with consistent exposure times.

### Total RNA extraction and quantitative real-time PCR (qPCR)

Total RNA was extracted from cultured cells or tissue samples using TRIzol reagent (Invitrogen, #15596026), followed by purification with a Direct-zol RNA Miniprep Kit (Zymo Research, #R2052). RNA integrity was assessed, and complementary DNA (cDNA) was synthesized from 1 µg of total RNA using the Transcriptor First Strand cDNA Synthesis Kit (Roche, #04896866001). Quantitative real-time PCR was performed on an ABI StepOne Plus system using SYBR Green Master Mix (Roche, #04707516001). Each reaction was run in technical triplicates, and all experiments included at least three independent biological replicates. The mRNA expression levels of the target genes were normalized to those of an internal reference gene, and relative quantification was performed using the comparative ΔΔCT method.

### Cell culture and transfection

Human embryonic kidney 293T (HEK293T) cells were obtained from FuHeng Cell Center (Shanghai). Jurkat (Clone E6-1), RAW264.7 cell lines were acquired from YiZeFeng Biotechnology (Shanghai). All mammalian cell lines were maintained at 37°C in a 5% CO_2_ atmosphere. HEK293T, RAW264.7 cells were cultured in Dulbecco’s modified Eagle’s medium (DMEM; Gibco #10569010) supplemented with 10% fetal bovine serum (FBS; Gibco #16000044) and 1% penicillin‒streptomycin (100×). Jurkat cells were grown in RPMI-1640 medium supplemented with 10% FBS and 1% penicillin‒streptomycin. HEK293T cells were transfected with polyethylenimine (PEI; Polysciences #24765-1). *Drosophila melanogaster* Schneider’s S2 cells were cultured in Sf-900™ II SFM (Gibco #10902088) supplemented with 50 U/mL penicillin and 50 µg/mL streptomycin and incubated at 25°C. S2 cells were transfected at 60–75% confluency using a DDAB suspension (250 µg/mL). All the cell lines used in this study are listed in Table EV16.

### Western blotting

Protein samples were lysed in ice-cold buffer containing 50 mM Tris-Cl (pH 7.4), 1% Triton X-100, 150 mM NaCl, 1 mM EDTA, and a protease inhibitor cocktail. The lysates were subsequently centrifuged at 20,000 × g at 4°C for 30 minutes to remove insoluble debris. The resulting supernatants were mixed with 5× protein loading buffer and denatured for subsequent immunoblotting analysis. The following primary antibodies were used at the indicated dilutions: mouse anti-actin (Cell Signaling Technology, 1:1,000), mouse anti-Flag (Sigma‒Aldrich, 1:1,000), and mouse anti-HA (Cell Signaling Technology, 1:1,000). Horseradish peroxidase (HRP)-conjugated anti-rabbit or anti-mouse secondary antibodies (Jackson ImmunoResearch Laboratories) were used at a dilution of 1:10,000. Protein bands were visualized using an enhanced chemiluminescence (ECL) detection system (Pierce) according to the manufacturer’s instructions.

### Coimmunoprecipitation (Co-IP) in HEK293T cells

To detect protein‒protein interactions, HEK293T cells were seeded in 10 cm culture plates and transfected at approximately 70–80% confluency. Each plate was transfected with 8 µg of each plasmid DNA using 48 µl of polyethylenimine (PEI; Polysciences, Inc., #24765-1) at a concentration of 1 µg/µl. Forty-eight hours after transfection, the cells were lysed on ice with cell lysis buffer (25 mM Tris-HCl (pH 7.6), 150 mM NaCl, 10 mM MgCl_2_, 1% NP-40, and EDTA-free protease inhibitor cocktail). The lysates were briefly sonicated for 15 seconds and clarified by centrifugation at 20,000 × g at 4°C for 20 min. A total of 2 mg of protein from the supernatant was incubated with anti-HA (Thermo Scientific, #88837) or anti-c-MYC (Thermo Scientific, #88842) magnetic beads at 4°C for 2–3 h with hours of rotation. Bead rotation. The samples were subsequently washed three times with TBS (50 mM Tris-HCl, 150 mM NaCl, pH 7.4). Bound proteins were eluted by boiling in 1× SDS loading buffer containing 5% 2-mercaptoethanol for 5 min, after which the eluates were analyzed by western blotting.

### Bimolecular Fluorescence Complementation (BiFC) Assay

The Bimolecular Fluorescence Complementation (BiFC) assay was employed to directly visualize protein‒protein interactions in living cells. In this study, the coding sequences of the two proteins of interest were cloned and inserted into the pcDNA3.1-mVenusN and pcDNA3.1-mVenusC vectors, which encode the N-terminal (VN) and C-terminal (VC) fragments of mVenus, respectively. The constructed plasmids were cotransfected into HEK293T cells using the polyethylenimine (PEI) method. After 48 hours of incubation, reconstituted fluorescence was observed using a confocal laser scanning microscope. A positive protein‒protein interaction was indicated by the presence of a yellow fluorescent signal in the transfected cells.

### Protein expression and purification

The cDNA of *C. elegans* KIN-1 was inserted into a modified pGEX vector to express an N-terminal GST-tagged fusion protein. The cDNA of *C. elegans* AST-1 was cloned and inserted into a modified pET28a vector, yielding a construct with an N-terminal His6 tag. Standard PCR-based mutagenesis was employed when applicable. All the constructs were verified by DNA sequencing. Recombinant KIN-1 and AST-1 proteins were expressed in *Escherichia coli* BL21(DE3) cells at 16°C. The cells were harvested and resuspended in lysis buffer. KIN-1 was purified using Glutathione Sepharose 4B (Cytiva) affinity chromatography, while AST-1 was purified via Ni^2+^-Sepharose 6 Fast Flow (GE Healthcare) affinity chromatography. The purified proteins were dialyzed against buffer containing 25 mM HEPES (pH 7.5) and 150 mM NaCl. Finally, the proteins were concentrated, aliquoted, and stored at –80°C.

### In vitro phosphorylation assay

Phosphorylation reactions were carried out in a 20 µL volume containing kinase assay buffer (25 mM HEPES, pH 7.4; 150 mM NaCl; 10 mM MgCl_2_; 1 mM DTT), 1 µg of purified kinase, 2 µg of substrate protein, and 100 µM ATP. The reaction mixture was incubated at 30°C for 30 minutes to allow phosphorylation to proceed. Reactions were terminated by boiling the samples with 1×SDS loading buffer for 5 minutes. The denatured proteins were then resolved by SDS‒PAGE. For Pro-Q Diamond staining, the gel was first fixed in a solution containing 50% methanol and 10% acetic acid for 30 minutes, followed by three 10-minute washes with distilled water. The gel was then stained with Pro-Q Diamond phosphoprotein gel stain (Invitrogen, #P33301) for 90 minutes according to the manufacturer’s protocol. Destaining was subsequently performed using a buffer composed of 20% acetonitrile and 50 mM sodium acetate (pH 4.0) three times for 30 minutes each. After a final 10-minute wash with distilled water, the gel was scanned under UV irradiation to visualize phosphoprotein signals.

### Yeast One-Hybrid Assay

The DNA bait sequence was subsequently cloned and inserted into the pAbAi vector to generate the pBait-AbAi plasmid, which was subsequently integrated into the Y1HGold yeast genome to construct the bait/reporter strain. The background aureobasidin A (AbA) resistance of the Y1HGold bait strain was assessed by plating 10 µL of yeast culture onto SD/-Ura agar medium supplemented with varying concentrations of AbA (ranging from 100 to 1000 ng/mL). The minimal AbA concentration that completely suppressed the growth of the bait strain—or a slightly higher concentration—was selected for screening the binding proteins. The cDNA encoding the transcription factor of interest was cloned and inserted into the pGADT7 vector and transformed into the bait strain. Subsequently, 10 µL of serial dilutions (10⁻¹, 10⁻², and 10⁻³) of the transformed yeast were spotted onto SD/-His selection medium supplemented with 300 ng/mL AbA, followed by incubation at 30°C for 3 days. Both positive and negative controls were included in all the experiments.

### Luciferase Reporter Assay

For luciferase reporter assays, approximately 2×10^5^ 293T cells were seeded per well in a 24-well plate 24 hours prior to transfection. Cells were cotransfected with 500 ng of the pGL3-Pro firefly luciferase reporter plasmid and 1 ng of the pSV40-seapansy Renilla luciferase internal control plasmid. After 48 hours of incubation, the cells were harvested, and luciferase activity was measured using a dual-luciferase assay system (Promega, #E1960) according to the manufacturer’s instructions. Firefly luciferase activity was normalized to Renilla luciferase activity for each sample. All transfections were performed in three independent biological replicates.

### Electrophoretic mobility shift assay

An electrophoretic mobility shift assay (EMSA) was conducted to analyze DNA‒protein interactions using a LightShift Chemiluminescent EMSA Kit (Thermo Scientific, #20148) following the manufacturer’s instructions. Biotin 5′-end-labeled DNA probes were generated by PCR amplification with specific primers; the biotin-labeled promoter sequences used in this study are listed in the table below. Recombinant His-tagged proteins were expressed and purified as previously described. For supershift assays, 1 µL of anti-His antibody was added per reaction to confirm the specificity of the DNA‒protein complex.

### RNA interference and lentivirus production

The targeted shRNA sequences were subsequently cloned and inserted into the pGREEN-puro plasmid. For gene knockdown, cells were transfected with shRNA plasmids by either electroporation or lentiviral transduction. Cells were harvested 48 hours post-transfection for subsequent functional analyses. For recombinant protein expression, the cDNA sequences encoding TTR52, LAT1, and CED1 were cloned and inserted into plasmid 113891 to generate C-terminal GFP-Flag-tagged fusion constructs (TTR52–GFP-Flag, LAT1–GFP, CED1–GFP), with GFP alone serving as a control. Each plasmid was individually packaged into lentivirus in 293TN cells, as described above. The resulting viruses were used to transduce HEK293T cells. Stably transduced cells were selected, and the expressed fusion proteins were purified using anti-Flag magnetic bead–based immunoprecipitation.

For lentivirus production, 293TN cells were cotransfected with the shRNA-expressing pGREEN-puro plasmid and the lentiviral packaging plasmids pSBH155 and pSBH156 using polyethylenimine (PEI). After 24 hours, the transfection medium was replaced with fresh DMEM supplemented with 10% fetal bovine serum (FBS) and 1% penicillin‒streptomycin. Viral supernatants were collected at 48 and 72 hours post-transfection, pooled, and used as the lentiviral stock for subsequent infections.

### Lipid dot-blot assay

Lipid‒protein interactions were analyzed using prespotted membrane lipid strips (P-6002; Echelon Biosciences). The strips were initially blocked in Tris-buffered saline containing 0.1% Tween-20 (TBST) with 3% bovine serum albumin (BSA) for 1 hour at room temperature. The membranes were ubsequently incubated with 2 nM target protein in blocking buffer at 4°C for 6 hours with gentle agitation. After incubation, the strips were washed three times with TBST and probed with an anti-FLAG primary antibody (1:1,000 dilution) for 1 hour at room temperature. The membranes were subsequently incubated with an HRP-conjugated secondary antibody and visualized using an enhanced chemiluminescence (ECL) detection system.

### RNA interference in S2 cells

Primers for dsRNA synthesis were designed using the online Drosophila RNAi screening platform (https://fgr.hms.harvard.edu/fly cell-based-rnai). A T7 promoter sequence (TAATACGACTCACTATAGGG) was appended to the 5′ end of each primer. Double-stranded RNA (dsRNA) was synthesized using the High Yield RNA Synthesis Kit (NEB), with the corresponding PCR products used as templates. All sequences of primers used are listed in Table EV3. For RNAi treatment, 500 µl of S2 cells were incubated with 5 µg of dsRNA for 36 hours. FITC-labeled apoptotic cells (ACs) were subsequently added and incubated at 25°C for 12 h to allow efferocytosis. The cells were then collected for qPCR analysis and microscopic examination.

### Apoptosis, effferocytosis and degradation assays in S2 cells

Apoptosis induction was performed as described previously (Nagano, Ui-Tei et al., 2000). S2 cells were treated with 0.25 μg/mL actinomycin D (AcD; MCE#HY-17559) at 25°C for 18 h, at which time apoptosis was induced, and the surface of the dying cells were exposed to PS. Apoptotic cells were added to fresh S2 cells at a ratio of 10:1, allowed to undergo efferocytosis for 6 h, and then analyzed by confocal microscopy. For RNAi treatment of apoptotic cells, efferocytosis assays were performed as described previously (Yamashita, Suzuki et al., 2020). In brief, S2 cells were incubated with 5 μM CellTracker™ Red CMTPX (Thermo Fisher #C34552) at 25°C for 1 h and then treated with 0.25 μg/mL AcD at 25°C for 1 h. The cells were subsequently collected by centrifugation at 1000 × g for 3 min and suspended in Sf-900TM II SFM. CMTPX-labeled prey cells (3×105 cells) were incubated with 1×105 S2 cells at 25°C for 60 min and analyzed by confocal microscopy on glass-bottom 96-well plates (Thermo Fisher Scientific). For the degradation assay in S2 cells, a stable cell line, CharON, which expresses a caspase-activated GFP and a pH-insensitive mCherry, was treated as described above, added to S2 cells after apoptosis induction at a ratio of 10:1, and incubated for 6 hours. As the lysosomes are acidified to allow the degradation of apoptotic cells, which quenches the green fluorescence signal, we analyzed the degradation efficiency using the remaining GFP fluorescence signal.

### Nuclear and cytoplasmic protein extraction

Cells were harvested and washed with ice-cold phosphate-buffered saline (PBS) to remove cellular debris and contaminants. After centrifugation (e.g., 500 × g, 5 min at 4°C), the supernatant was carefully aspirated, and the cell pellet was retained. For cytoplasmic protein extraction, the pellet was resuspended in cold lysis buffer containing protease inhibitors. The suspension was vortexed thoroughly and incubated on ice for 10–15 minutes to lyse the plasma membrane while keeping the nuclear membrane intact. Prolonged lysis was avoided to prevent the leakage of nuclear proteins. The lysate was then subjected to high-speed centrifugation at 12,000 × g for 5 minutes at 4°C. The resulting supernatant, containing cytoplasmic proteins, was collected and stored at –80°C. The pellet, enriched in nuclei, was washed with wash buffer to remove residual cytoplasmic contamination, followed by another round of centrifugation. For nuclear protein extraction, the purified nuclear pellet was resuspended in nuclear extraction buffer. The sample was vigorously vortexed or sonicated to disrupt the nuclear membrane, followed by incubation on ice for 30 minutes with intermittent mixing. After centrifugation at 12,000 × g for 5 minutes at 4°C, the supernatant containing nuclear proteins was collected and stored at –80°C for downstream applications. All procedures were performed on ice or at 4°C to maintain protein stability and subcellular compartment integrity.

### Mice

*Adgrl3*-deficient mice were generated on a C57BL/6JJY background using a conventional knockout approach. The *Adgrl3* knockout allele was introduced by gene targeting, and heterozygous (*adgrl3 ^*+* /^* ^*−*^) mice were generated (C57BL/6JCya-Adgrl3em1/Cya). Heterozygous mice were bred to produce offspring for genotyping and further colony expansion. Homozygous *Adgrl3 ^*−* / *−*^* mice were generated by intercrossing *Adgrl3^*+*/*−*^* litters. Genotyping was performed by PCR using tail genomic DNA with specific primer sets (F1/R1 and F2/R1) to distinguish wild-type (*^+/+^*), heterozygous (^*+*/*−*^), and homozygous (*^*−*/*−*^*) genotypes. Mice carrying a targeted deletion of 1425 bp in the Adgrl3 gene were confirmed by PCR analysis and Sanger sequencing. The experimental mouse strains were kept in the Animal Experiment Center of Shaanxi Normal University according to standard methods.

### Generation of murine bone marrow-derived macrophages (BMDMs)

Murine bone marrow-derived macrophages (BMDMs) were generated as previously described, with minor modifications. Briefly, bone marrow (BM) cells were isolated from the femurs and tibiae of mice by flushing with cold phosphate-buffered saline (PBS). Red blood cells were lysed using ACK lysis buffer, and the nucleated cells were collected by centrifugation. Cells were resuspended in complete Dulbecco’s modified Eagle’s medium (DMEM), which consisted of DMEM basal medium (Fisher Scientific, 11-995-073) supplemented with 10% heat-inactivated fetal bovine serum (HI-FBS; Gibco, A5256801), 2 mM L-glutamine (Fisher Scientific, 25-030-081), and 20% L-929 cell-conditioned medium as a source of macrophage colony-stimulating factor (M-CSF). BM cells were seeded at a density of 2 × 10^6^ to 3 × 10^6^ cells per well in 6-well nontissue culture-treated plates and cultured at 37°C in a humidified atmosphere containing 5% CO_2_. The differentiation medium was replaced with fresh medium on days 3 and 4. After 7 days of culture, the adherent cells exhibited a typical macrophage morphology and were used as mature BMDMs for subsequent in vitro assays.

### Phagocytosis Assay of Apoptotic Cells

To induce apoptosis, CharON-expressing Jurkat cells were exposed to ultraviolet (UV) radiation at a dose of 150 J/m^2^ and subsequently incubated for 3 h at 37°C under 5% CO_2_ to allow apoptosis progression. The resulting apoptotic Jurkat cells were harvested, resuspended in sterile 1× PBS, and added to RAW264.7 cells or bone marrow-derived macrophages (BMDMs) previously transfected with target-specific shRNA at a ratio of 5:1 (apoptotic cells: macrophages).

For time-lapse confocal imaging, macrophages were cocultured with UV-induced apoptotic CharON-expressing Jurkat cells that carried a fluorescent reporter gene. After 2 h of coincubation, noninternalized apoptotic cells were gently removed by washing the macrophages twice with PBS. For microscopy analysis, 1 mL of DMEM was added to each culture dish for observation. Efferocytosis was quantified as the percentage of macrophages containing internalized red fluorescent CharON Jurkat cells among the total number of macrophages per field of view (FOV). Degradation was assessed as the ratio of macrophages containing internalized CharON Jurkat cells exhibiting only red fluorescence (indicative of loss of the green signal due to lysosomal degradation) to those containing cells with both red and green fluorescence, expressed as a percentage per field of view.

The coculture period for flow cytometry analysis was 2 h. Subsequently, the macrophages were rinsed twice with PBS to remove noninternalized cells, detached using trypsin, resuspended in 500 μL of PBS, and analyzed using a CytoFLEX S flow cytometer (Beckman Coulter). The data were processed using CytExpert software. Consistent with these findings, compared with control macrophages, macrophages with impaired degradation function exhibited significantly greater TAMRA fluorescence intensity.

### Measurement of Intracellular cAMP Levels

Intracellular cyclic adenosine monophosphate (cAMP) concentrations were quantified using the E-EL-0056 cAMP Competitive ELISA Kit (Elabscience Biotechnology Co., Ltd.) following the manufacturer’s standard protocol with minor adaptations for cell-based samples. Briefly, cells transfected with control shRNA, *adgrl3* shRNA, or *adgrl3* shRNA plus rescue constructs (wild-type ADGRL3 or the ΔGPS mutant) were washed twice with ice-cold PBS and then lysed in ice-cold 0.1 M HCl containing 0.1% Triton X-100. Lysates were incubated on ice for 10 min to inhibit endogenous phosphodiesterase activity and centrifuged at 12,000 × g for 10 min at 4°C to collect the clear supernatants. The total protein concentration of each sample was determined using the BCA assay to normalize cAMP levels across groups. All samples were processed according to the manufacturer’s instructions, including the acetylation step for enhanced sensitivity, and absorbance was measured at 450 nm using a microplate reader. cAMP concentrations were calculated from the standard curve, normalized to the total protein content, and expressed as relative fold changes compared to the control shRNA group. All assays were performed in triplicate with three independent biological replicates each. Statistical differences between groups were analyzed using one-way ANOVA followed by Tukey’s post-hoc test, with p < 0.05 considered statistically significant.

### Statistical analyses

Statistical analyses were performed using Microsoft Excel, and bar graphs or other statistical plots were generated using GraphPad Prism software. The data were obtained from at least three biological replicates. Details of the statistical tests, sample sizes, and methodologies are provided in the individual figure legends. The data are presented as the mean ± SEM. One-way analysis of variance (ANOVA) or Student’s t test was used to determine the statistical significance. “*” represents p < 0.05, “**” represents p < 0.01, “***” represents p < 0.001.

## Data Availability

All study data are included in the article and/or supporting information.

## Acknowledgments

This work was partially supported by the National Natural Science Foundation of China

(Grant No. 32370799 to Hui Xiao), the National Natural Science Foundation of China (Grant No. 31871387 to Hui Xiao), the National Natural Science Foundation of China Youth Program (Grant No. 32300622 to Lei Yuan), the Sanqin Bochuang Talent Support Program of Shaanxi Province (Grant No. 2024SQBC004 to Lei Yuan), and the Innovative Research Team for the Central Universities (Grant No. GK202302003 to Hui Xiao).

## Author Contributions

**Fuqiao Liu**: Data curation, Software, Methodology, Formal analysis, Writing – original draft. **Lei Yuan:** Data curation, Software, Methodology, Formal analysis. **Qian Zheng:** Data curation, Software, Methodology, Formal analysis. **Yang Kang**: Data curation, Software, Methodology. **Aowei Wang**: Data curation, Software, Methodology. **Kuo Li**: Data curation, Software, Methodology. **Yunmin Xie:** Data curation, Software, Methodology. **Lu Chen**: Data curation, Software, Methodology. **Peiyao Li:** Data curation, Software, Methodology. **Hui Wang**: Validation, Investigation, Visualization. **Zhi Li**: Data curation, formal analysis, resources, validation, conceptualization, writing–review and editing. **Hui Xiao**: Conceptualization, Resources, Data curation, Formal analysis, Supervision, Funding acquisition, Validation, Investigation, Visualization, Methodology, Writing – original draft, Writing – review and editing.

## Competing interests

The authors declare that they have no competing interests.

---

**Fig. EV1.**
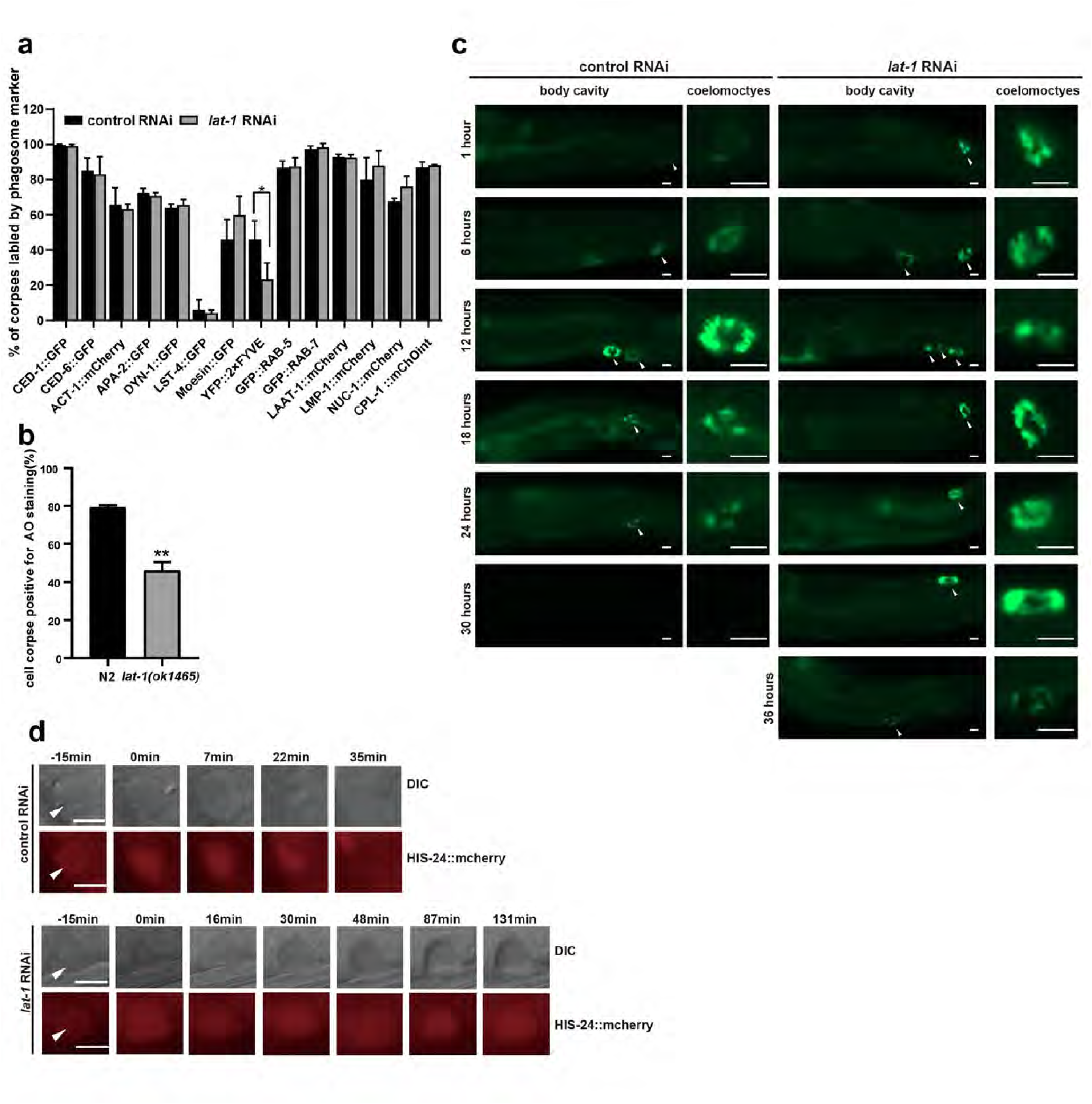
LAT-1 is required for phagolysosomal maturation. (a) Quantification (mean ± SEM) of cell corpses labeled with different phagosomal markers. A total of ≥100 cell corpses were analyzed for each marker and data were derived from three replicates. Comparisons were performed between the N2/control and *lat-1* RNAi worms for each marker. (b) Quantification (mean ± SEM) of AO-positive germ cell corpses in N2 and *lat-1* mutants. (c) The uptake and degradation of ssGFP in coelomocytes were monitored at the indicated time points in arIs36/control and *lat-1* RNAi worms (heat shocked for 60 min at 33°C). Left, accumulation of ssGFP in the body cavity (400×); arrows indicate coelomocytes that are enlarged in the right pictures (1000×). Bars, 10 μm. (d) Time-lapse chasing of HIS-24::mCherry positive phagolysosomes in N2/control and *lat-1* RNAi worm germlines. The time point at which HIS-24::mCherry mostly rings the cell corpse was set to 0 min. Arrows indicate continuous presence of cell corpses (DIC, top row) and HIS-24::mCherry (fluorescence, bottom row). DIC and fluorescence images were obtained at certain time intervals. Bars, 5 µm. The unpaired *t* test was performed. *P < 0.05, **P < 0.01. All error bars indicate the mean ± SEM.

**Fig. EV2.**
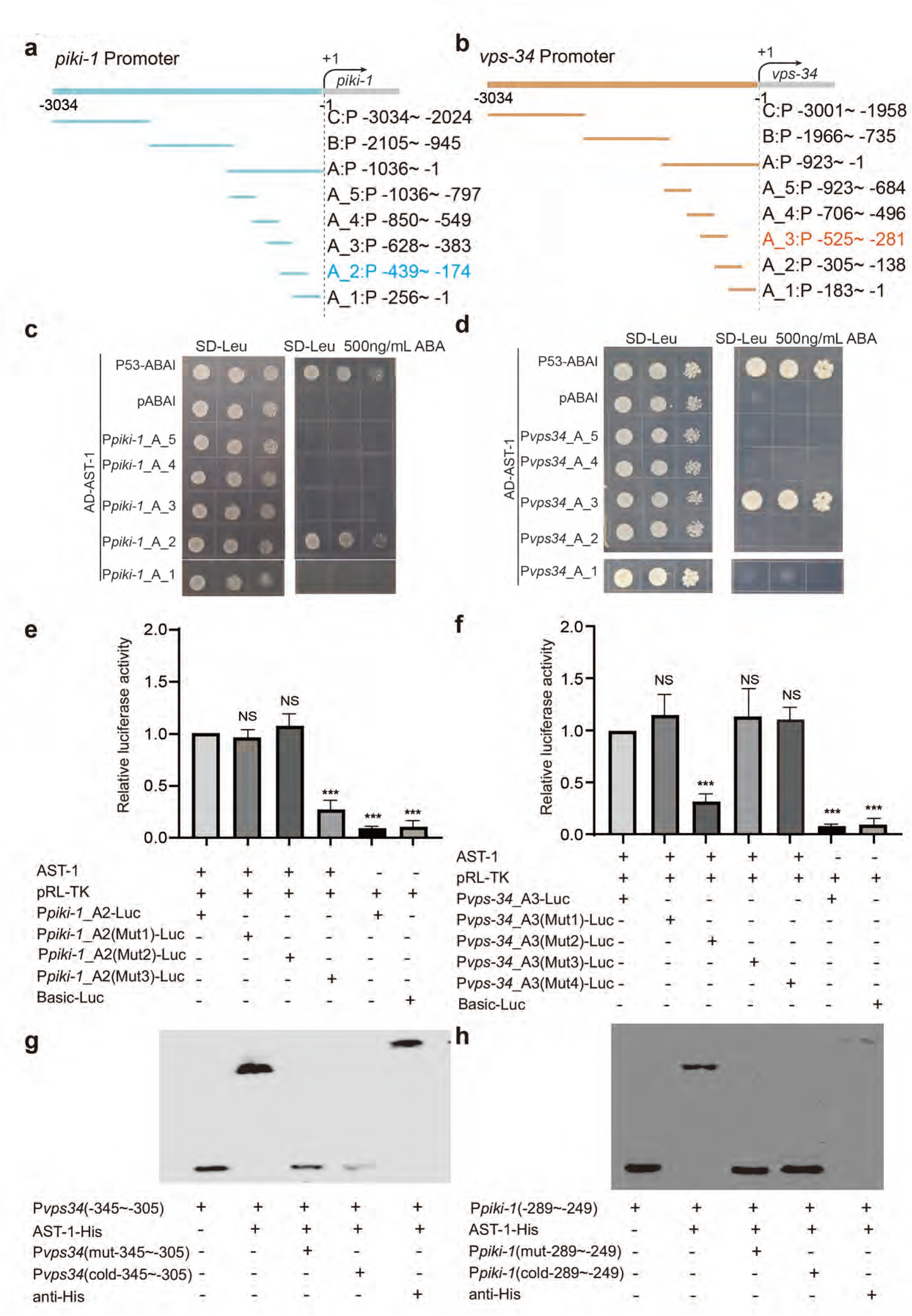
AST-1 regulates the transcription of *vps-34* and *piki-1* by directly binding to its promoter. (a-b) Schematic diagrams of the segmented promoter regions of *piki-1* (a) and *vps-34* (b). (c-d) Yeast one-hybrid assays testing the binding of AST-1 to the corresponding promoter regions of *vps-34* (c) and *piki-1* (d) as defined in (a) and (b). (e-f) Luciferase reporter assays evaluating the activity of *vps-34* (e) and *piki-1* (f) promoters in response to AST-1 using wild-type and site-directed mutant promoter constructs. The mean activity with the standard deviation of three independent transfections is represented. Statistical significance was assessed using one-way ANOVA. (g-h) Electrophoretic mobility shift assays (EMSA) showing specific binding of purified AST-1::His to biotin-labeled promoter probes of *vps-34* (g) and *piki-1* (h). The assays included supershift with anti**-**His antibody and competition with unlabeled wild**-**type or mutant probes.

**Fig. EV3.**
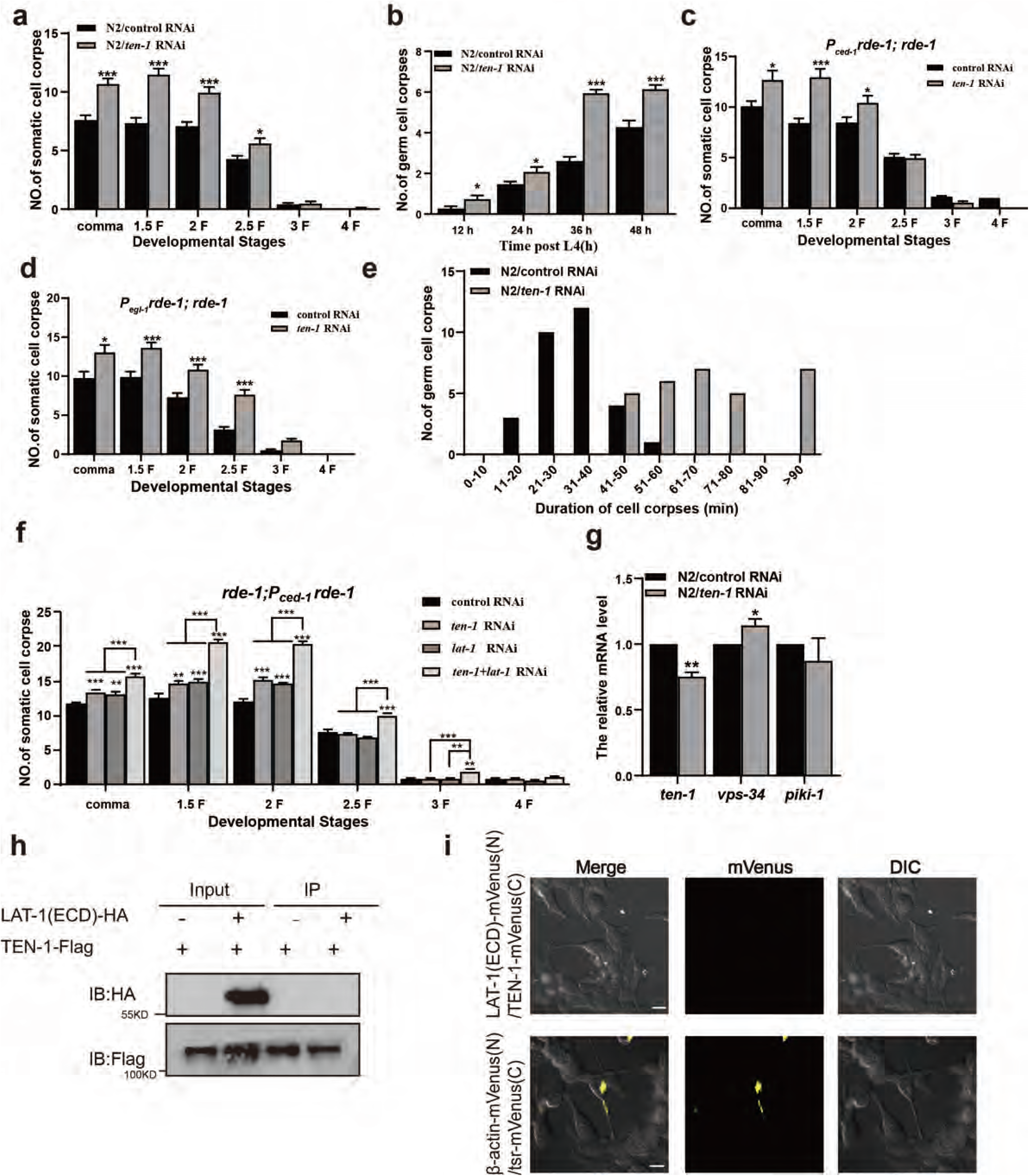
*ten-1* does not function in the same genetic pathway with *lat-1* for cell corpse removal. (a-d and f) Embryo development of (a, c, d, and f) and adult (h post L4) (b) stages was quantified (mean ± SEM) for the indicated strains. Fifteen embryos and adult worms were scored at each stage for each strain. (e) Four-dimensional microscopy analysis of germ cell corpse duration in N2/control and *ten-1* RNAi worms. The persistence of 30 germ cell corpses was monitored in each group. (g) Relative mRNA levels of *vps-34* and *piki-1* in N2/control and *ten-1* RNAi worms were determined using qRT-PCR. (h) Co-IP assays in 293T cells to assess the interaction between LAT-1 (ECD) (tagged with HA) and TEN-1 (Flag-tagged). Proteins were immunoprecipitated using anti-HA magnetic beads. (i) BiFC assay detecting the interaction between the extracellular domains of LAT-1 and TEN-1. Scale bar, 10 μm.

**Fig. EV4.**
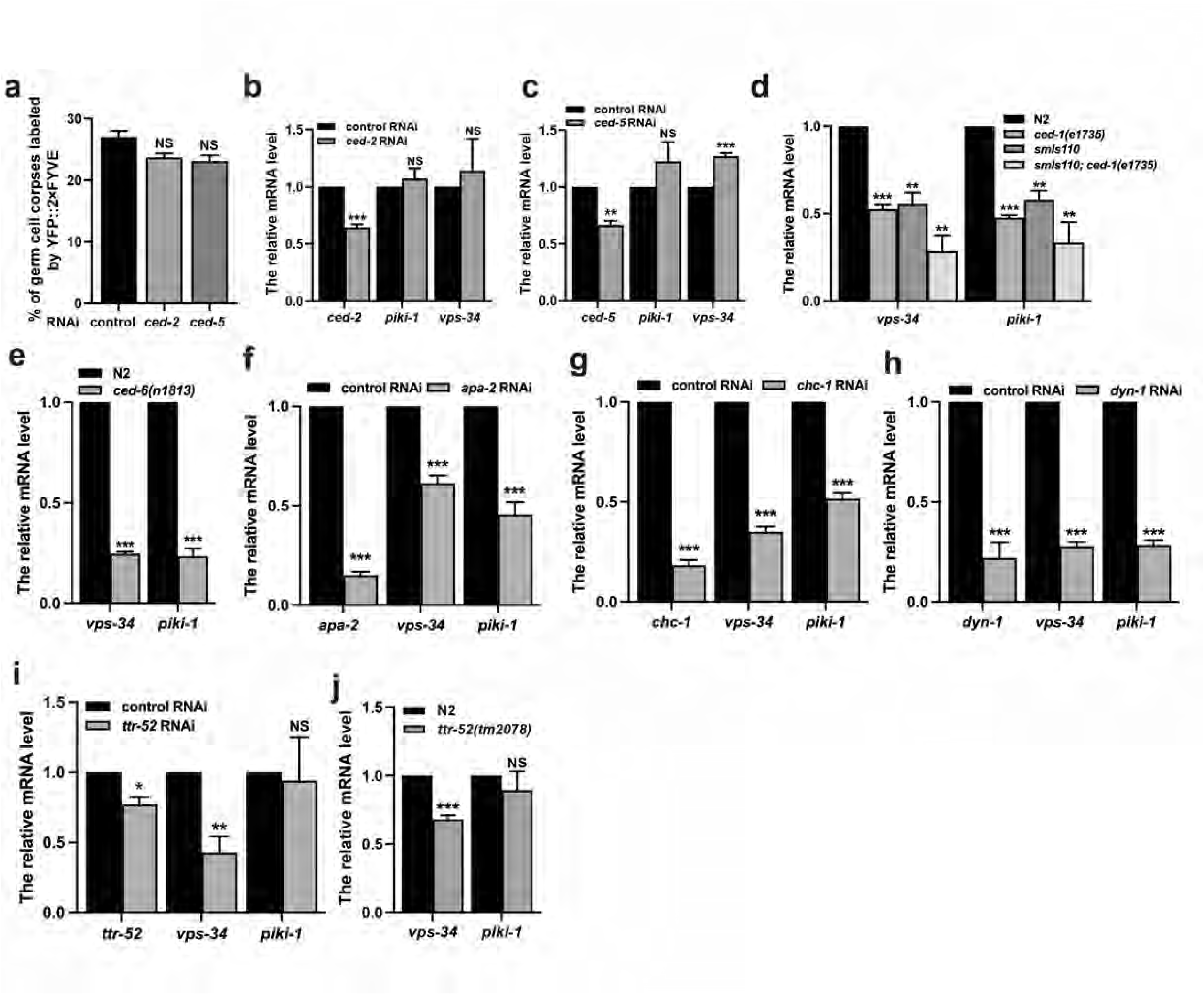
The intracellular domain of CED-1 is essential for its full function in regulating *vps-34* and *piki-1* transcription. (a) Quantification (mean ± SEM) of cell corpse labeling by the phagosomal marker YFP::2×FYVE. A total of ≥100 cell corpses were analyzed, and the data were derived from three independent replicates for each experiment. Comparisons were performed between control and *ced-2* or *ced-5* RNAi worms using the YFP::2×FYVE marker. (b-j) Relative mRNA levels of *vps-34* and *piki-1* in N2/control and *ced-2* RNAi (b), N2/control and *ced-5* RNAi (c), N2, *ced-1(e1735)*, and overexpression strains (d), N2 and *ced-6(n1813)* (e), N2/control and *apa-2* RNAi (f), N2/control and *chc-1* RNAi (g), N2/control and *dyn-1* RNAi (h), N2/control and *ttr-52* RNAi (i), and N2 and *ttr-52(tm2078)* (j). The unpaired t test was performed. *P < 0.05, **P < 0.01, ***P < 0.001. All error bars indicate the mean ± SEM.

**Fig. EV5.**
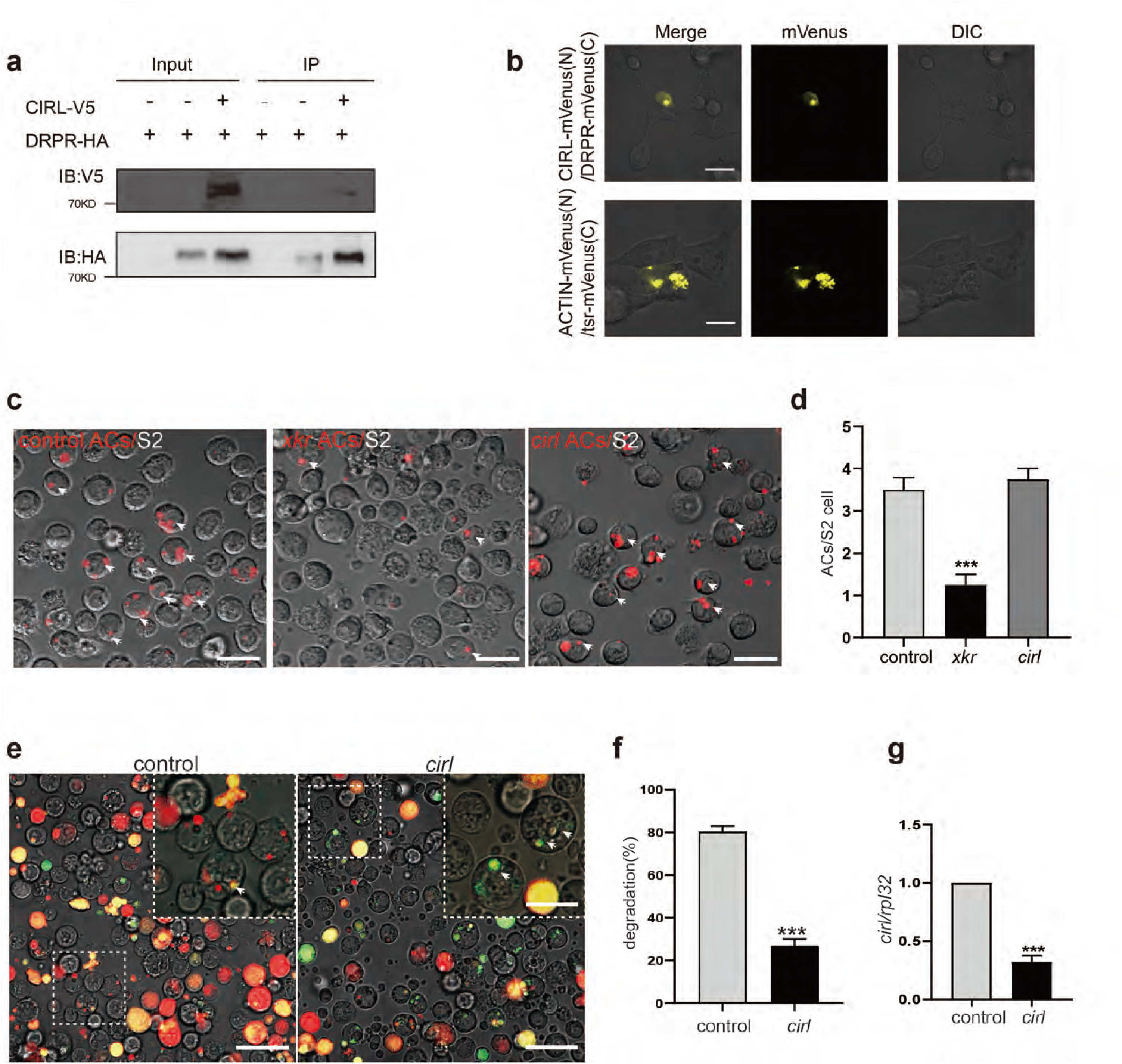
*cirl* is required for apoptotic cell degradation in drosophila S2 cells. (a) Co-IP assays in S2 cells to assess the interaction between DRPR (tagged with HA) and CIRL (tagged with V5). Proteins were immunoprecipitated using anti-HA-magnetic beads. (b) BiFC assay detecting the interaction between DRPR and CIRL. Scale bar, 10 μm. (c). In vitro efferocytosis. S2 cells, *xkr^ko^* cells (positive control), and cirl RNAi cells labeled with CellTracker Red were treated with 0.25 μg/mL AcD for 1h, and then added to fresh S2 cells for 1h of incubation. Red particles (ingested apoptotic cells, ACs) were observed and counted using confocal fluorescence microscopy. Three independent experiments were conducted. The regions enclosed by the rectangles are shown at a higher magnification in the insets. Scale bars, 20 μm. (d). The column diagram shows the statistical results of the ACs per S2 cell in (c). *** *p* < 0.001 (one-way ANOVA). (e). Degradation assay in S2 cells. The CharON sTable EV2 cell line, which expressed a caspase-activated GFP and a pH-insensitive mCHERRY, was treated with 0.25 μg/mL actinomycin D for 6h, and then added to control RNAi cells and cirl RNAi cells, respectively, and observed by confocal fluorescence microscopy. (f). The column diagram shows the statistical results of the single red fluorescence signal of phagocytes in (e). *** *p* < 0.001 (Student’s t test). (g). The mRNA level of cirl after RNAi in S2 cells was detected by qRT-PCR, and rpl32 was used as the internal control. Three independent experiments were performed. (Student’s t test).

**Fig. EV6.**
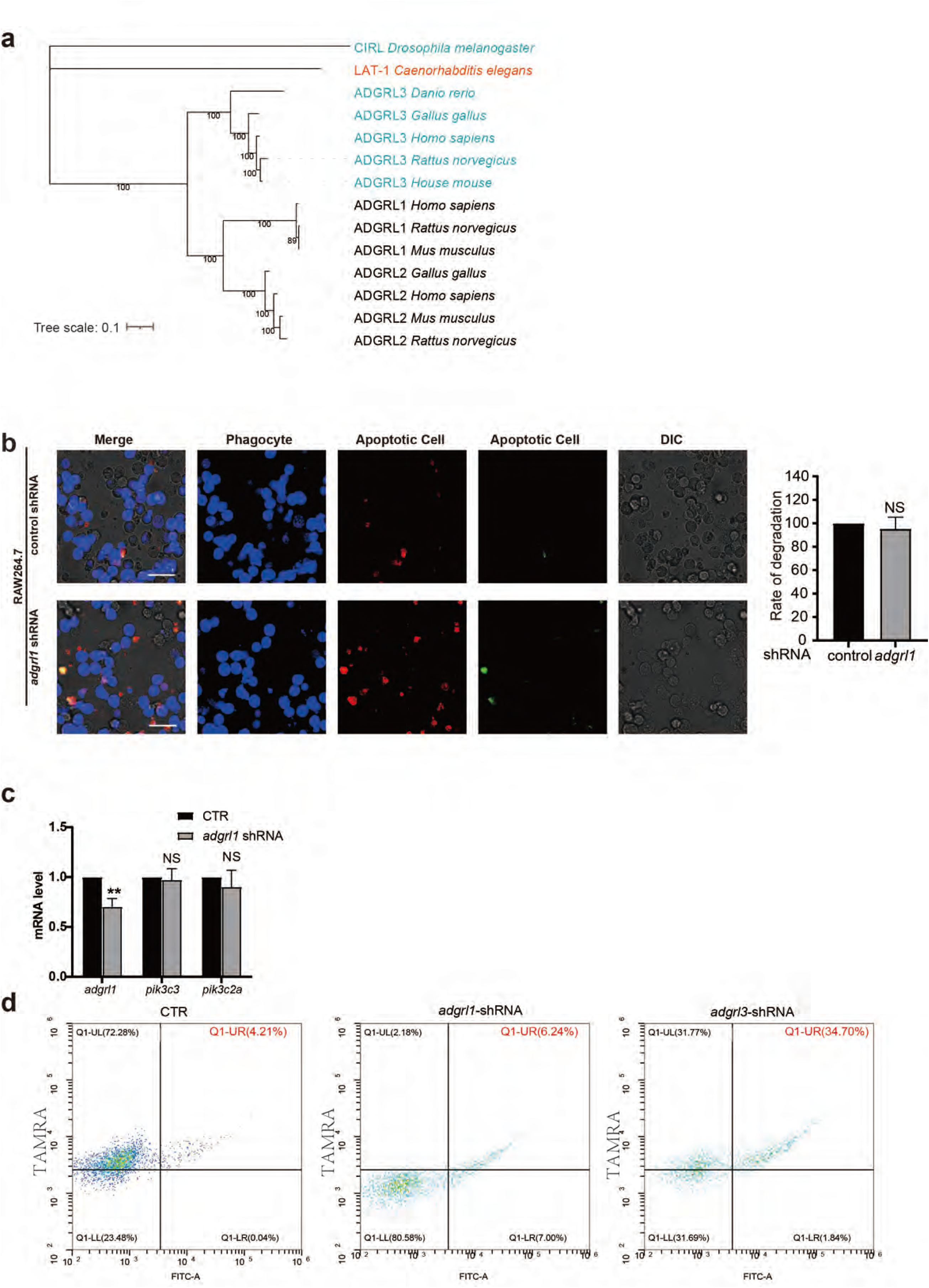
*adgrl1* is not involved in efferocytosis. (a) Phylogenetic tree of LAT-1 orthologs constructed using the maximum likelihood method across representative species. (b) The role of *adgrl1* in macrophage degradation of apoptotic cells. Macrophages expressing a control or *adgrl1*-targeting shRNA (blue) were incubated with dual-color apoptotic CharON Jurkat reporters (green: apoptosis-induced, acid-sensitive; red: acid-resistant). Student’s t test was used for statistical analysis. Relative levels were quantified and presented as mean ± SEM. Data are representative of at least three independent experiments. For each experiment, five random imaging fields were analyzed. *P < 0.05, **P < 0.01, ***P < 0.001; NS, not significant. Scale bars, 10 μm. (c) Effect of *adgrl1* knockdown on the relative mRNA levels of *pik3c3* and *pik3c2a*. Student’s t test was used for statistical analysis. Relative levels were quantified and presented as mean ± SEM. Data are representative of at least three independent experiments. *P < 0.05, **P < 0.01, ***P < 0.001; NS, not significant. (d) Flow cytometric analysis of efferocytosis following *adgrl1* or *adgrl3* knockdown. Macrophages with knockdown of the indicated genes were co-cultured with TAMRA-labeled apoptotic cells, and efferocytosis was quantified using flow cytometry.

**Fig. EV7.**
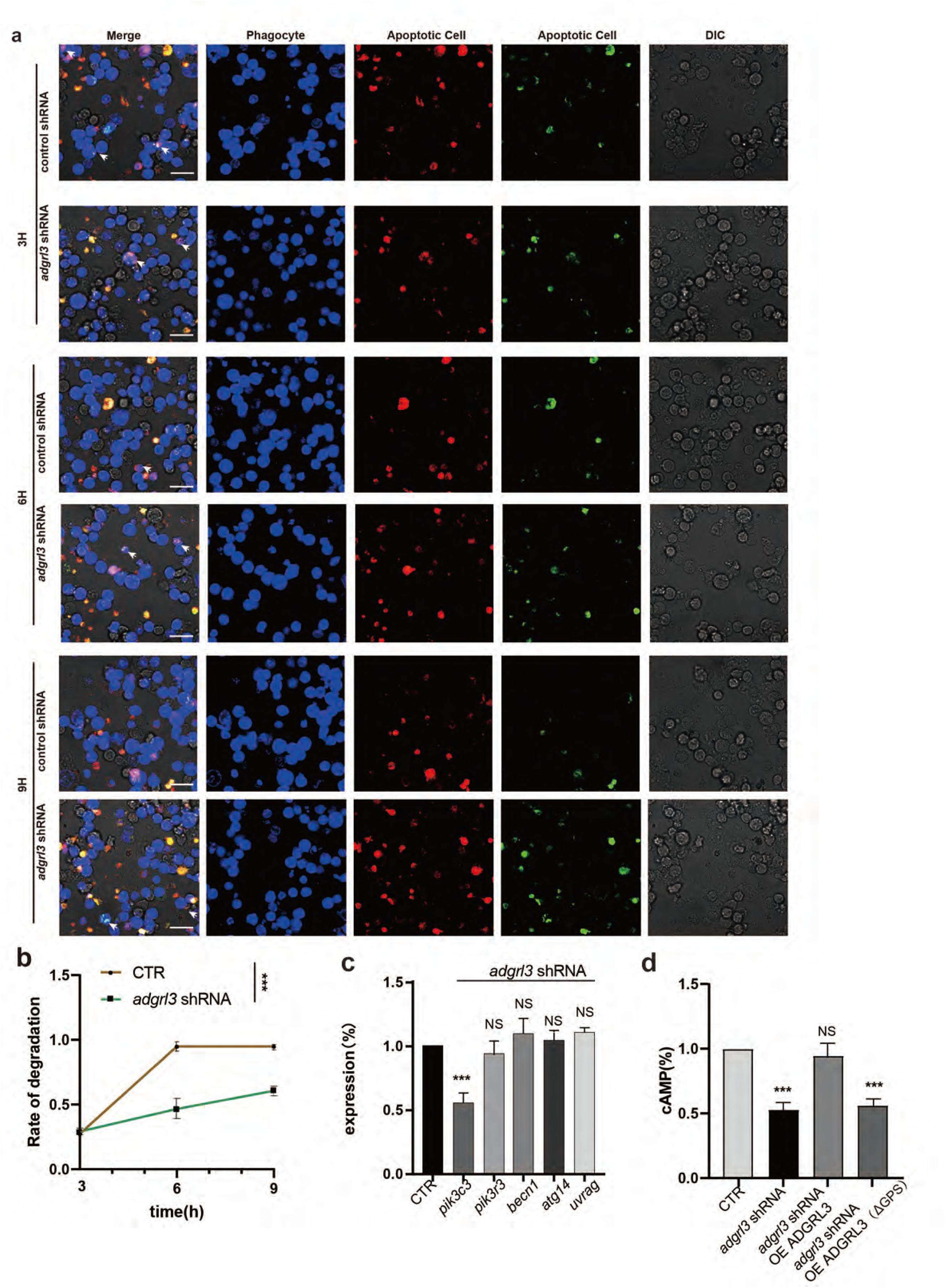
*adgrl3* regulates apoptotic cell degradation. (a) Role of *adgrl3* in macrophage degradation of apoptotic cells. Macrophages expressing control or *adgrl3*-targeting shRNA (blue) were incubated with dual-color apoptotic CharON Jurkat reporters (green: apoptosis-induced and acid-sensitive; red: acid-resistant). Data are representative of at least three independent experiments. Student’s t test was used for statistical analysis. Relative levels were quantified and presented as mean ± SEM. For each experiment, five random imaging fields were analyzed. *P < 0.05, **P < 0.01, ***P < 0.001; NS, not significant. Scale bars, 10 μm. (b) Quantification of the effect of *adgrl3* knockdown on apoptotic cell degradation. Data are representative of at least three independent experiments performed. Five random imaging fields were analyzed for each experiment. *P < 0.05, **P < 0.01, ***P < 0.001; NS, not significant. (c) Effect of *adgrl3* knockdown on the relative mRNA levels of *pik3c3* and *pik3c2a*. Student’s t test was used for statistical analysis.Data are representative of at least three independent experiments. *P < 0.05, **P < 0.01, ***P < 0.001; NS, not significant. (d) Analysis of intracellular cAMP levels. Cells were subjected to adgrl3 knockdown and subsequent rescue with wild-type ADGRL3 or the ΔGPS mutant. cAMP was quantified using the E-EL-0056 ELISA kit. Data are shown as the mean ± SEM of three biological replicates. Data are representative of at least three independent experiments performed. *P < 0.05, **P < 0.01, ***P < 0.001; NS, not significant.

**Fig. EV8.**
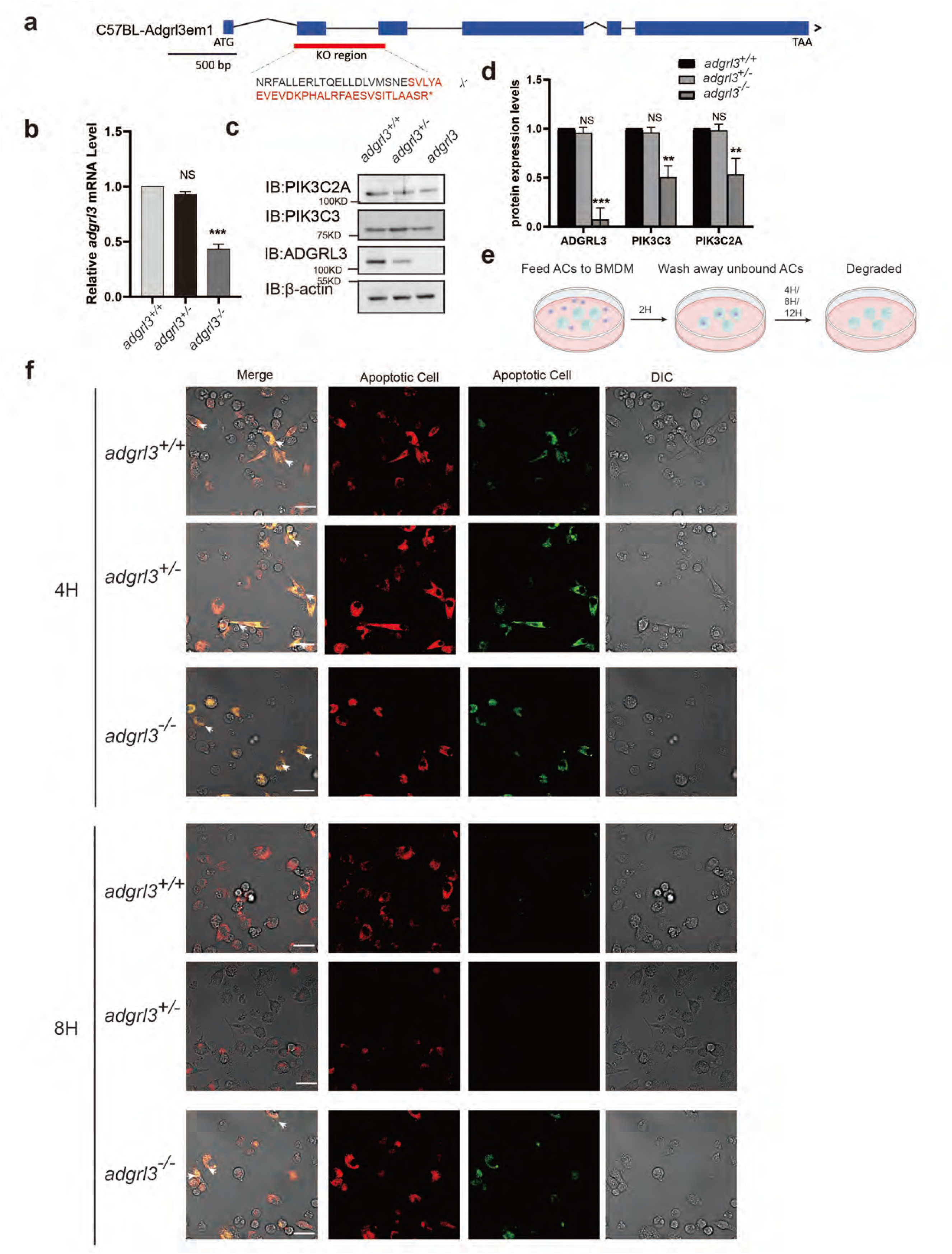
Role of ADGRL3 in the degradation of apoptotic cells by BMDMs. (a) Diagram of ADGRL3 protein and targeted deletion strategy. (B-D) adgrl3 mRNA expression levels in control(*adgrl3*^+/+^), ADGRL3 heterozygous(*adgrl3*^⁺/⁻^), and ADGRL3 KO(*adgrl3*^⁻/⁻^) BMDMs. (c-d) Analysis of the related protein levels. Western blot detection of ADGRL3, PIK3C3, and PIK3C2A in control(*adgrl3*^+/+^), ADGRL3 heterozygous(*adgrl3*^⁺/⁻^), and ADGRL3 KO (*adgrl3*^⁻/⁻^) BMDMs. (e) Experimental workflow for assessing the effect of ADGRL3 on phagocytosis and degradation of apoptotic cells by BMDMs. (f) Role of *adgrl3* in degradation of apoptotic cells in BMDMs. Macrophages expressing a control or *adgrl3*-targeting shRNA (blue) were incubated with dual-color apoptotic CharON Jurkat reporters (green: apoptosis-induced, acid-sensitive; red: acid-resistant). One-way ANOVA followed by post-hoc test was used. Data are representative of at least three independent experiments. For each experiment, five random imaging fields were analyzed. *P < 0.05, **P < 0.01, ***P < 0.001; NS, not significant. Scale bars, 10 μm.

**Fig. EV9.**
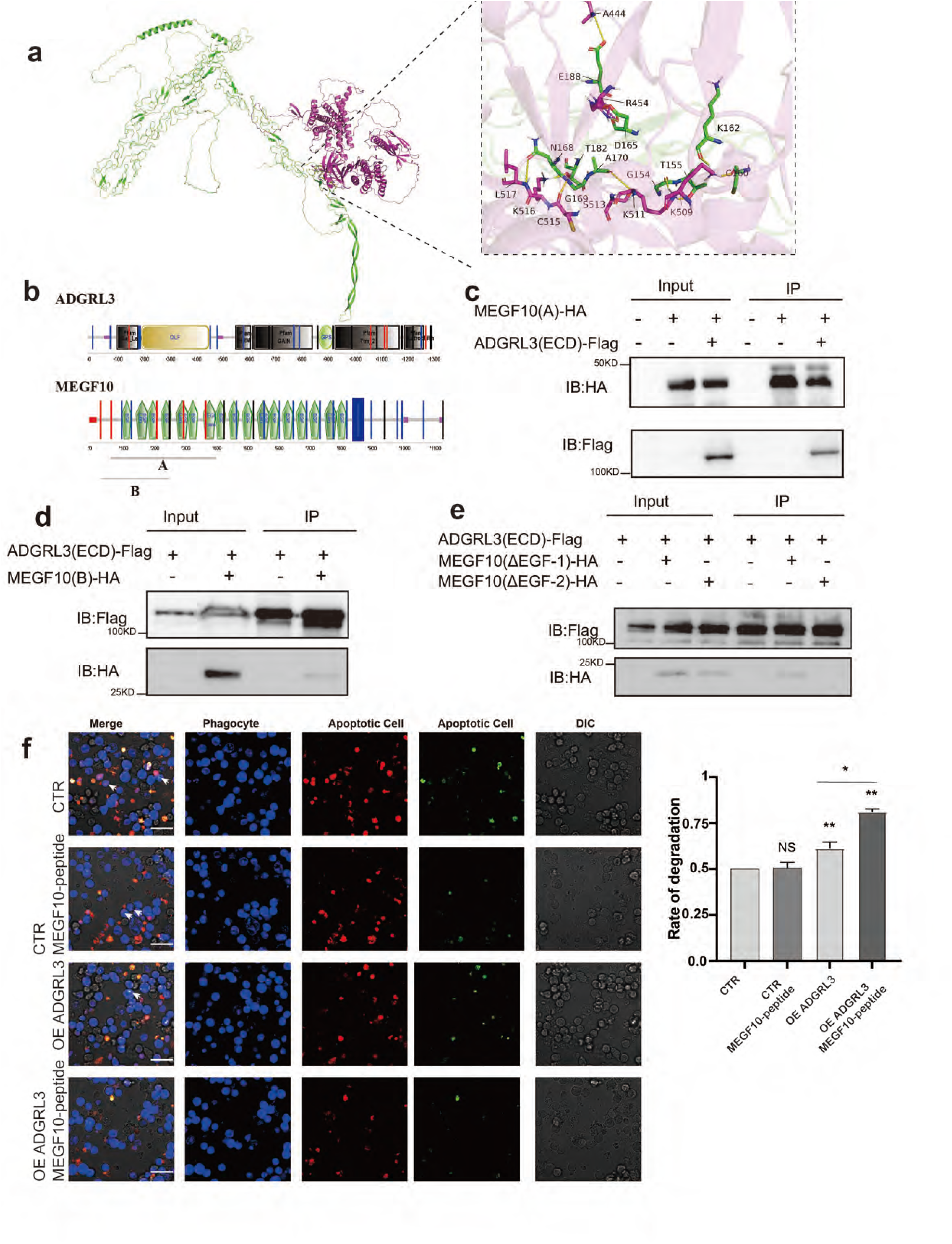
The MEGF10–ADGRL3 Interaction Enhances Efferocytosis through a Conserved EGF-like Domain. (a) The three-dimensional structure of the MEGF10-ADGRL3 protein complex was predicted using AlphaFold2. The left panel shows the overall structural model, with MEGF10 colored in green and ADGRL3 in purple; the right panel presents a magnified view of the key interaction interface. (b) Domain composition and functional domain prediction analysis of MEGF10 and ADGRL3. (c-e) Co-immunoprecipitation (Co-IP) assays were performed to examine the interaction capabilities of various truncated mutants of MEGF10 with its binding proteins. (f) Functional validation experiments were conducted in RAW264.7 macrophages: ADGRL3 was overexpressed, and an exogenous peptide containing the EGF-like domain of MEGF10 was added. After 4 hours of treatment, phagocytic activity was assessed. One-way ANOVA followed by post-hoc test was used. Data are representative of at least three independent experiments. For each experiment, five random imaging fields were analyzed. *P < 0.05, **P < 0.01, ***P < 0.001; NS, not significant. Scale bars, 10 μm.

**Fig. EV10.**
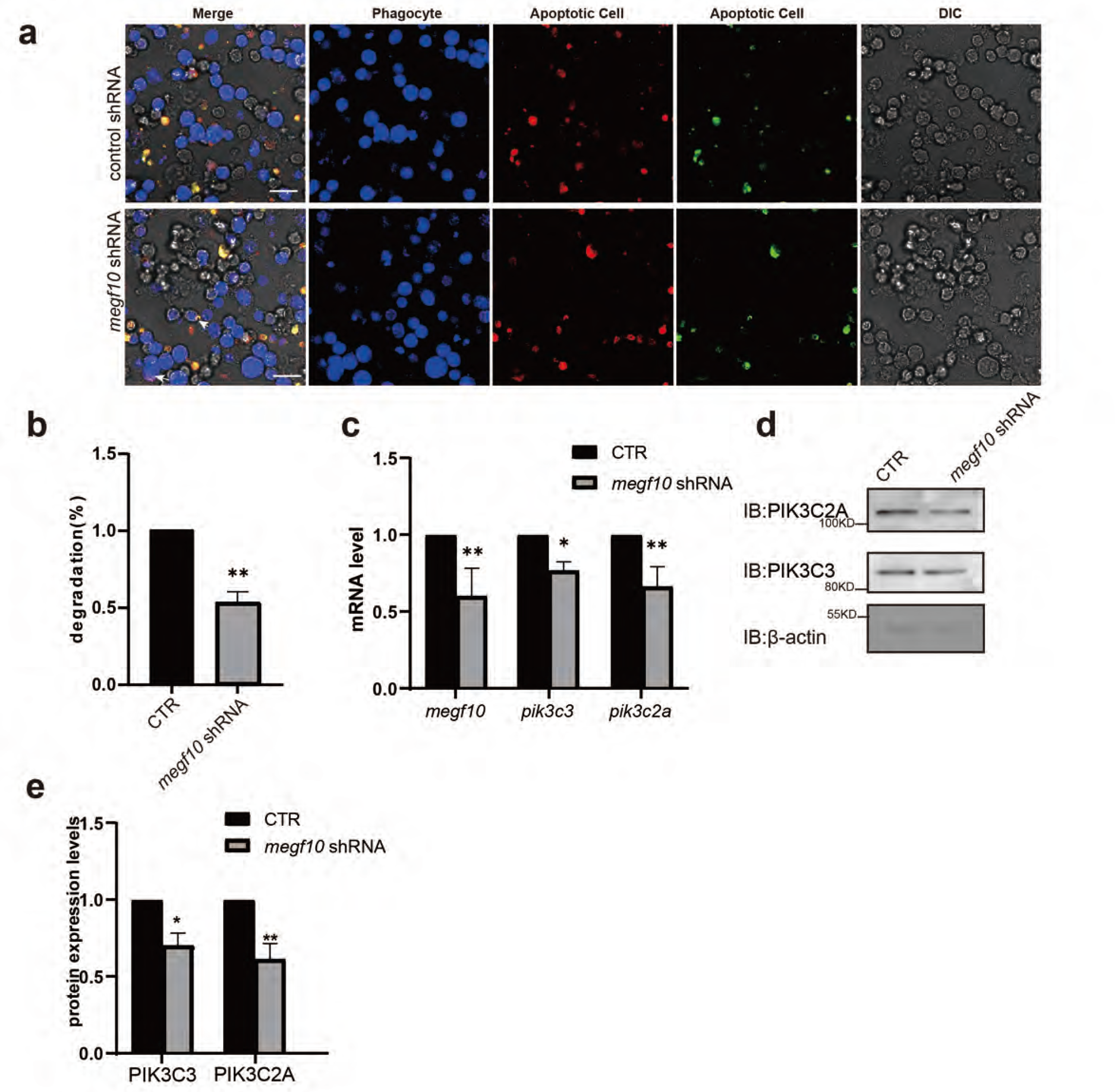
*megf10* regulates apoptotic cell degradation. (a) Role of *megf10* in macrophage degradation of apoptotic cells. Macrophages expressing control or *megf10*-targeting shRNA (blue) were incubated with dual-color apoptotic CharON Jurkat reporters (green: apoptosis-induced and acid-sensitive; red: acid-resistant). Scale bars, 10 μm. (b) Student’s t test was used for statistical analysis. Quantification of the effect of *megf10* knockdown on apoptotic cell degradation. The data are representative of at least three independent experiments. Five random imaging fields were analyzed for each experiment. *P < 0.05, **P < 0.01, ***P < 0.001; NS, not significant. (c) Effect of *megf10* knockdown on the relative mRNA levels of *pik3c3* and *pik3c2a*. Student’s t test was used for statistical analysis. Data are representative of at least three independent experiments. *P < 0.05, **P < 0.01, ***P < 0.001; NS, not significant. (d) Effect of *megf10* knockdown on the relative protein levels of PIK3C3 and PIK3C2A. (e) Quantitative analysis of relative PIK3C3 and PIK3C2A protein levels based on western blot band intensities in panel (d). Student’s t test was used for statistical analysis. Data are representative of at least three independent experiments. *P < 0.05, **P < 0.01, ***P < 0.001; NS, not significant.

**Fig. EV11.**
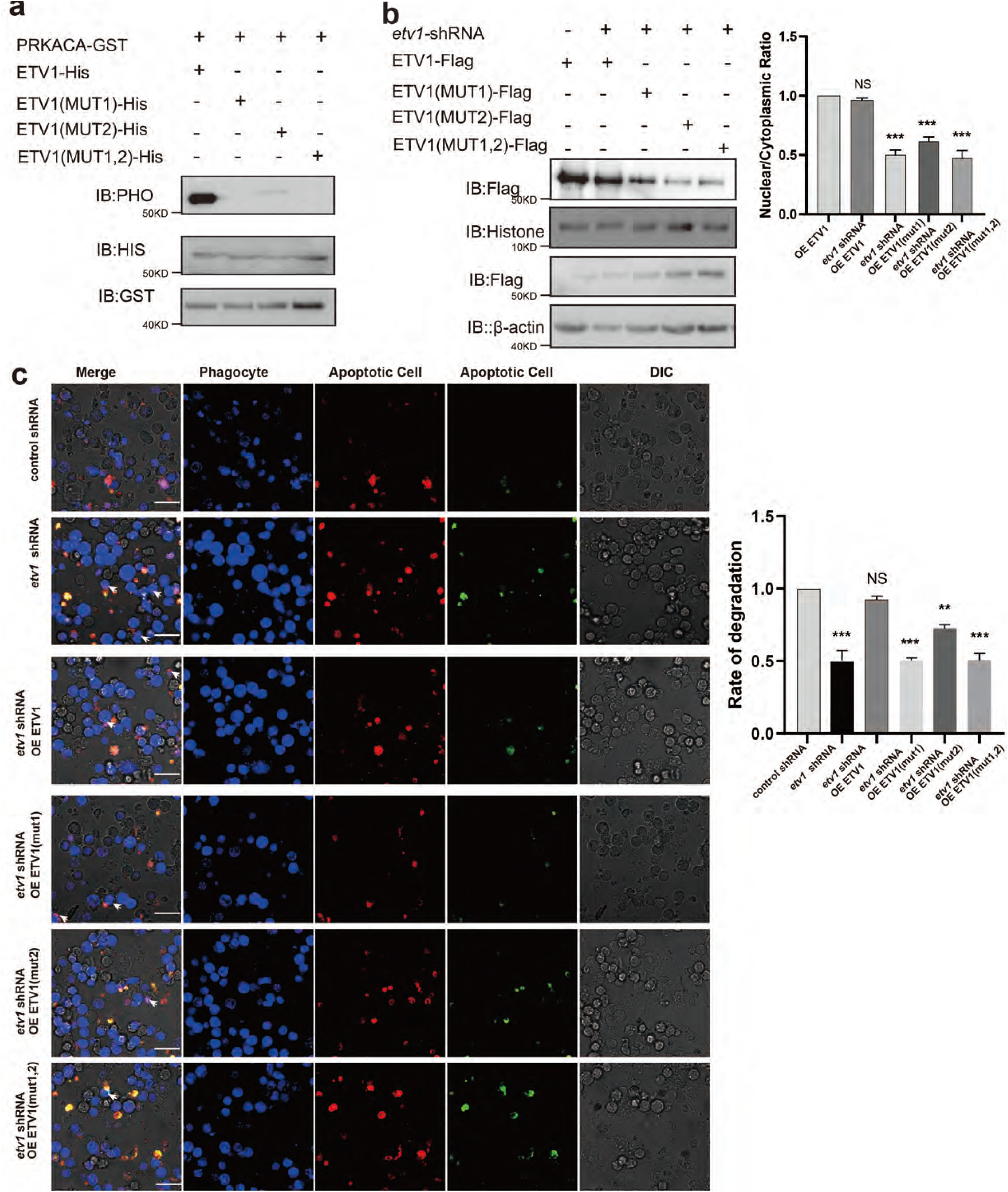
Functional and Molecular Characterization of ETV1 Phosphorylation Mutants in RAW264.7 Cells. (a) In vitro phosphorylation site identification. An in vitro kinase assay combined with Western blotting was employed to verify the direct phosphorylation of ETV1 by specific kinases and to precisely map the phosphorylation modification sites. (b)Nuclear translocation capacity assay of ETV1 mutants. Subcellular fractionation was performed to quantitatively compare the nuclear-to-cytoplasmic distribution ratios of wild-type and mutant ETV1, thereby assessing alterations in nuclear import capability. Experiments were independently repeated three times. (c)Functional validation of ETV1 phosphorylation sites. By constructing different single-point mutants of ETV1, the regulatory effects of each mutant on the degradation function of RAW264.7 cells were evaluated. Student’s t test was used for statistical analysis.Data are representative of at least three independent experiments. For each experiment, five random imaging fields were analyzed. *P < 0.05, **P < 0.01, ***P < 0.001; NS, not significant. Scale bars, 10 μm.

**Fig. EV12.**
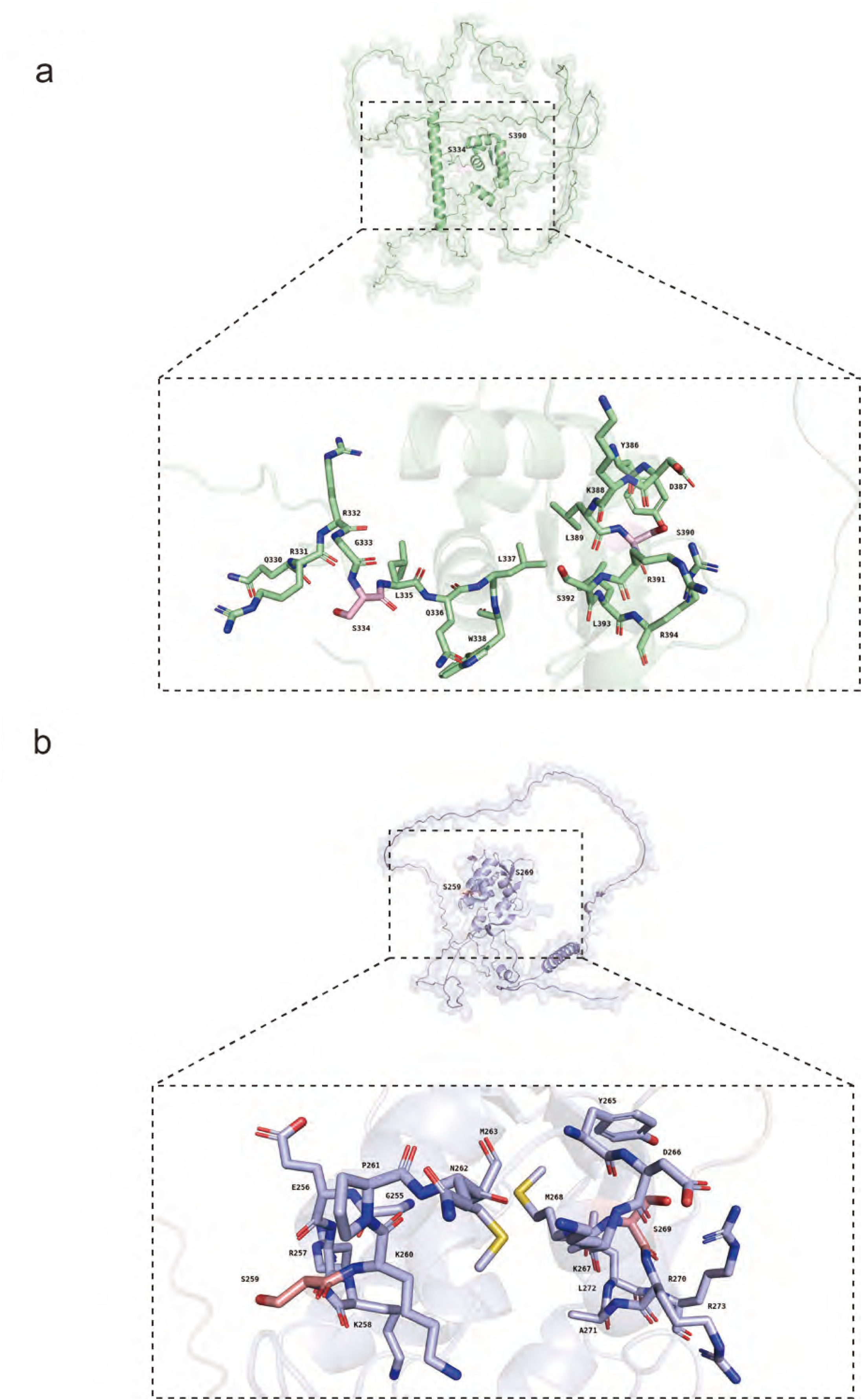
Prediction and evolutionary conservation analysis of PKA phosphorylation sites in ETV1 and AST-1. (a) Predicted three-dimensional structure of AST-1 and spatial distribution of PKA phosphorylation sites. (b) Predicted three-dimensional structure of ETV1 and corresponding mapping of phosphorylation sites.

**Fig. EV13.**
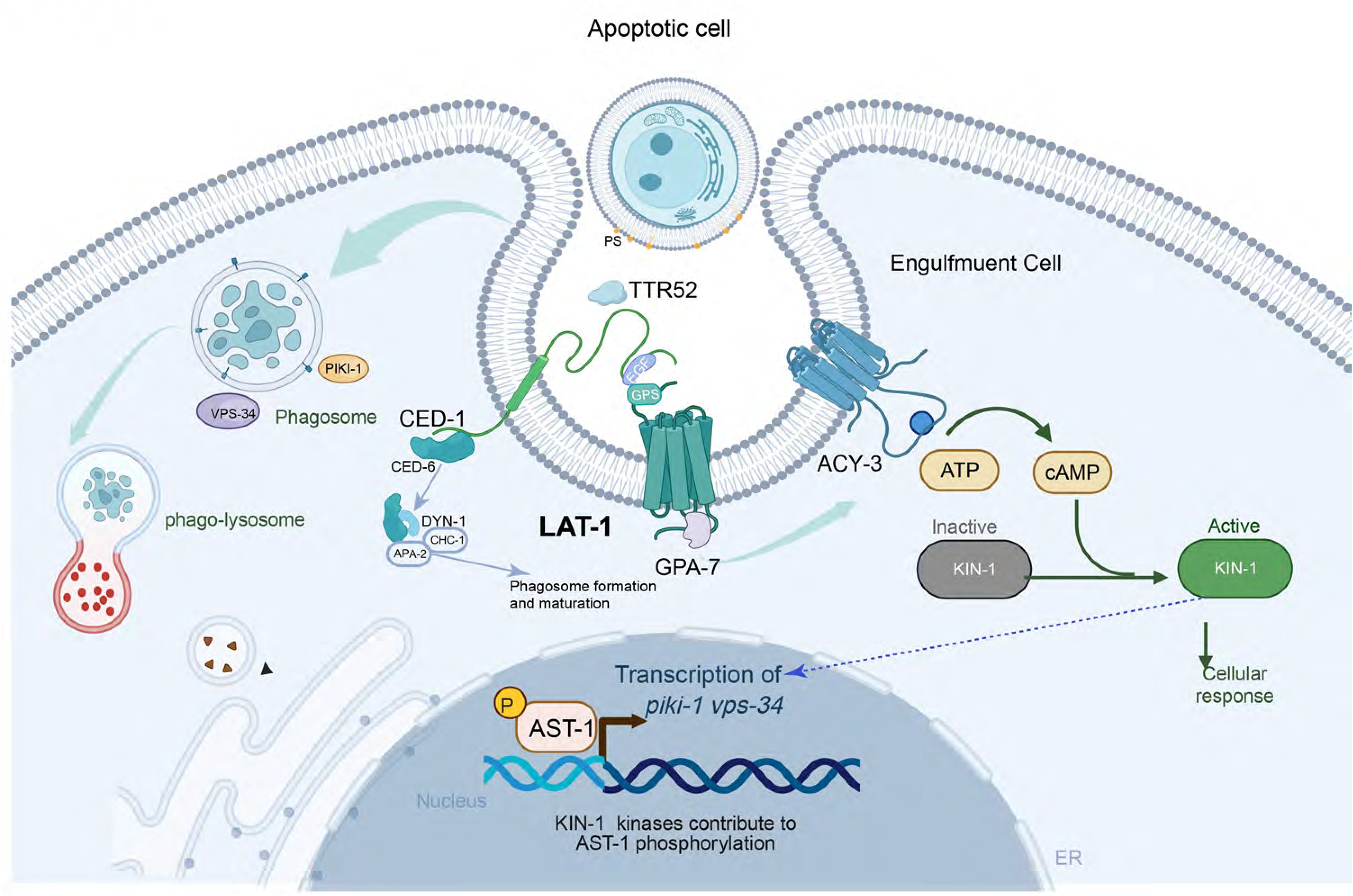
Pathway diagram annotationof LAT-1 function as a co-receptor for CED-1 in apoptotic cell degradation. During the engulfment of apoptotic cells, exposed phosphatidylserine (PS) on the dying cell surface is recognized by the engulfment receptor CED-1 in the engulfment cell, with a potential contribution from bridging molecules such as TTR52. LAT-1, an adhesion G protein-coupled receptor (aGPCR), acts as a co-receptor for CED-1 by interacting with the extracellular EGF-like repeats of the receptor via its GAIN domain, which facilitates LAT-1 activation. Activated LAT-1 couples to the G protein GPA-7, triggering a downstream signaling cascade. GPA-7 activates adenylyl cyclase ACY-3, which catalyzes the conversion of ATP to cyclic adenosine monophosphate (cAMP). Elevated intracellular cAMP levels activate protein kinase A (PKA), represented by the C. elegans ortholog KIN-1. Activated KIN-1 phosphorylates ETS-domain transcription factor AST-1. Phosphorylated AST-1 translocates to the nucleus, where it binds to the promoter regions of *piki-1* and *vps-34*, upregulating their transcription. Meanwhile, CED-1 signaling also initiates a parallel pathway via CED-6, DYN-1, APA-2, and CHC-1 to drive early phagosome formation and maturation. Transcriptionally upregulated PIKI-1, a class III phosphoinositide 3-kinase (PI3K), generates phosphatidylinositol-3-phosphate (PtdIns3P) during early phagosome sealing, whereas VPS-34 further enriches PtdIns3P on sealed phagosomes at later stages. Together, these two PI3 kinases promote phagosome maturation and phagosome–lysosome fusion, enabling the efficient degradation of apoptotic cell corpses. Thus, the LAT-1**→**GPA-7 **→** ACY-3 **→** cAMP **→** KIN-1 **→** AST-1 pathway converts engulfment signals into sustained gene expression changes, ensuring sufficient PI3K activity for robust phagosome maturation and apoptotic cell clearance.

